# Using three-dimensional regulatory chromatin interactions from adult and fetal cortex to interpret genetic results for psychiatric disorders and cognitive traits

**DOI:** 10.1101/406330

**Authors:** Paola Giusti-Rodríguez, Leina Lu, Yuchen Yang, Cheynna A Crowley, Xiaoxiao Liu, Ivan Juric, Joshua S Martin, Armen Abnousi, S. Colby Allred, NaEshia Ancalade, Nicholas J Bray, Gerome Breen, Julien Bryois, Cynthia M Bulik, James J Crowley, Jerry Guintivano, Philip R Jansen, George J Jurjus, Yan Li, Gouri Mahajan, Sarah Marzi, Jonathan Mill, Michael C O’Donovan, James C Overholser, Michael J Owen, Antonio F Pardiñas, Sirisha Pochareddy, Danielle Posthuma, Grazyna Rajkowska, Gabriel Santpere, Jeanne E Savage, Nenad Sestan, Yurae Shin, Craig A Stockmeier, James TR Walters, Shuyang Yao, Bipolar Disorder Working Group of the Psychiatric Genomics Consortium, Eating Disorders Working Group of the Psychiatric Genomics Consortium, Gregory E Crawford, Fulai Jin, Ming Hu, Yun Li, Patrick F Sullivan

**Affiliations:** University of North Carolina, Department of Genetics, Chapel Hill, NC, US; Case Western Reserve University, Department of Genetics and Genome Sciences, Cleveland, OH, US; University of North Carolina, Department of Biostatistics, Chapel Hill, NC, US; Cleveland Clinic, Department of Quantitative Health Sciences, Cleveland, OH, US; Cardiff University, MRC Centre for Neuropsychiatric Genetics and Genomics, Division of Psychological Medicine and Clinical Neurosciences, Cardiff, Wales, UK; King’s College London, Social, Genetic & Developmental Psychiatry Centre, Institute of Psychiatry, Psychology & Neuroscience, London, UK; South London and Maudsley Hospital, UK National Institute for Health Research Biomedical Research Centre, London, UK; Karolinska Institutet, Department of Medical Epidemiology and Biostatistics, Stockholm, Sweden; University of North Carolina, Department of Psychiatry, Chapel Hill, NC, US; Erasmus University Medical Center, Department of Child and Adolescent Psychiatry, Rotterdam, NL; Vrije Universiteit Amsterdam, Department of Complex Trait Genetics, Amsterdam, NL; Case Western Reserve University, Department of Psychiatry, Cleveland, OH, US; Louis Stokes VA Medical Center, Department of Psychiatry, Cleveland, OH, US; University of Mississippi Medical Center, Department of Psychiatry and Human Behavior, Jackson, MS, US; King’s College London, London, UK; University of Exeter, University of Exeter Medical School, Exeter, UK; Case Western Reserve University, Department of Psychological Sciences, Cleveland, OH, US; Yale School of Medicine, Department of Neuroscience and Kavli Institute for Neuroscience, New Haven, CT, US; Duke University, Center for Genomic and Computational Biology, Durham, NC, US; Duke University, Department of Pediatrics, Division of Medical Genetics, Durham, NC, US; Case Western Reserve University, Case Comprehensive Cancer Center, Cleveland, OH, US

## Abstract

Genome-wide association studies have identified hundreds of genetic associations for complex psychiatric disorders and cognitive traits. However, interpretation of most of these findings is complicated by the presence of many significant and highly correlated genetic variants located in non-coding regions. Here, we address this issue by creating a high-resolution map of the three-dimensional (3D) genome organization by applying Hi-C to adult and fetal brain cortex with concomitant RNA-seq, open chromatin (ATAC-seq), and ChIP-seq data (H3K27ac, H3K4me3, and CTCF). Extensive analyses established the quality, information content, and salience of these new Hi-C data. We used these data to connect 938 significant genetic loci for schizophrenia, intelligence, ADHD, alcohol dependence, Alzheimer’s disease, anorexia nervosa, autism spectrum disorder, bipolar disorder, major depression, and educational attainment to 8,595 genes (with 42.1% of these genes implicated more than once). We show that assigning genes to traits based on proximity provides a limited view of the complexity of GWAS findings and that gene set analyses based on functional genomic data provide an expanded view of the biological processes involved in the etiology of schizophrenia and other complex brain traits.

## Introduction

In the last decade, genome-wide searches for genetic variation fundamental to human maladies of exceptional public health importance became feasible ^1,2^. Genomic studies are particularly important for idiopathic psychiatric disorders that have few proven/reproducible biological risk factors despite twin-family studies conclusively establishing a role for inheritance ^3,4^. Hopes that protein-coding variation would provide a key to schizophrenia ^5,6^ have not eventuated as sizable exome sequencing studies have identified only two genes to date ^7,8^. In contrast, exome sequencing studies for autism identified ~100 genes with far fewer cases than for schizophrenia ^9^. The lack of exonic findings for schizophrenia is unfortunate given the range of available tools for experimental modeling of single genes but exonic findings are more prevalent for severe psychiatric syndromes with onset early in life.

Genome-wide association studies (GWAS) of common genetic variation have yielded more findings, particularly when the numbers of cases are large ^2,10^. A landmark 2014 GWAS for schizophrenia identified 108 significant loci ^11^ and a later study found 145 loci ^12^. Most of the “genetic architecture” of common psychiatric disorders and brain traits lie in common variants of relatively subtle effects identifiable by GWAS ^2^. Other complex human diseases have similar conclusions (e.g., type 2 diabetes mellitus) ^13^. There have been 3,029 GWAS publications that identified 31,976 significant associations for 2,520 diseases, disorders, traits, or lab measures (***URLs***, Q4 2018), and these are almost always common variants of subtle effects (median odds ratio, OR, 1.22) and ORs infrequently exceed 2 ^10^.

Although GWAS findings are surprisingly informative in aggregate across the genome ^14^-^16^, delivering strong hypotheses about their connections to specific genes has been challenging ^2^. Investigators often rely on genomic location to connect significant SNPs to genes but this is problematic as GWAS loci: usually contain many correlated and significant SNP associations over 100s of Kb, many genes expressed in tissues of interest, and long-range effects to genes far outside a locus ^17^-^19^. These issues have led to efforts to use brain functional architecture ^2^ to connect GWAS loci to specific genes using gene expression quantitative trait loci (eQTL), SNP prioritization algorithms, chromatin interactions, or other approaches ^12,17^-^21^. The lack of direct connections to genes constrains subsequent experimental modeling and development of improved therapeutics.

The 3D arrangement of chromatin in cell nuclei enables physical and regulatory interactions between genomic regions located far apart in 1D genomic distance ^22^. Chromatin conformation capture (3C) methods enable identification of 3D chromatin interactions *in vivo* ^23,24^ and can clarify GWAS findings. For example, an intergenic region associated with multiple cancers was shown to be an enhancer for *MYC* via a long-range chromatin loop ^25,26^, and intronic *FTO* variants are robustly associated with body mass but influence expression of distal genes via long-range interactions ^27^. To interpret GWAS results for psychiatric disorders, Roussos et al. ^20^ used 3C methods to identify an intergenic chromatin loop for *CACNA1C* (from intronic GWAS associations for schizophrenia and bipolar disorder to its promoter). Won et al. ^17^ and Wang et al. ^19^ used brain 3D chromatin interactome data to assert connections of some schizophrenia associations to specific genes.

These examples suggest that knowledge of the 3D chromatin interactome in human brain could help clarify the meaning of GWAS findings for psychiatric disorders, brain-based traits, and neurological conditions. We were particularly interested in schizophrenia ^12^ and intelligence ^28^ because these two traits have many shared loci, a significant negative genetic correlation ^12,28^, lower premorbid intelligence is a risk factor for schizophrenia ^29^, and intellectual disability is an important comorbidity of schizophrenia ^30^. Because 3D interactome data from human brain are limited, we sequenced adult and fetal cortical samples to the greatest depth to date to enable a detailed portrait of the brain 3D chromatin interactome. After establishing the quality and informativeness of these new brain Hi-C data, we compared proximity/location-based gene identification with those from high-confidence regulatory chromatin interactions for 10 brain traits with sizable GWAS. We found that location provided a limited view of the complexity of GWAS findings whereas connections based on functional data from human brain capture greater complexity. Gene set analyses based on functional genomic data provide an expanded view of the biological processes involved in schizophrenia and intelligence.

## Results

### General properties of chromatin organization in brain

In this section, we describe the brain Hi-C data we generated and compare them to external datasets to establish their relevance. We applied “easy Hi-C” (eHi-C) ^31^ to postmortem samples (*N*=3 adult temporal cortex and *N*=3 fetal cortex). Using eHi-C, we generated 10.4 billion reads and, following quality control, identified 1.323 billion high-confidence *cis*-contacts that enabled a 10 Kb resolution map of the chromatin interactome. To our knowledge, these are the deepest Hi-C data from human brain, and contain 22.5X as many *cis*-contacts for adult cortex and 1.56X as many for fetal cortex compared to the next largest datasets ^17,32^ (***Figure 1a***, ***Table S1***). We evaluated four Hi-C readouts: “A” (active) or “B” (inactive) compartments (100 Kb resolution) ^33^; topologically associated domains (TADs, 40 Kb resolution) segment the genome into Mb-sized regions within which chromatin interactions have a strong tendency to occur ^34,35^; frequently interacting regions (FIREs, 40 Kb resolution) are regions with significantly greater *cis*-connectivity than expected under a null hypothesis of random collisions ^32,36^ (and can occur as contiguous super FIREs ^37^); and chromatin interactions identify genomic regions that are physically proximal in the nuclear 3D space ^22,36^-^42^, and consisted of two 10 Kb anchors 20 Kb to 2 Mb apart. We generated or obtained RNA-seq, open chromatin (ATAC-seq), ChIP-seq (H3K27ac, H3K4me3, and CTCF), and eQTL data ^18,43,44^ from adult and fetal cortex (***Table S2***). By combining eHi-C, RNA-seq, ATAC-seq, and ChIP-seq data from adult and fetal brain, we identified high-confidence regulatory chromatin interactions (HCRCI). HCRCI had *P* < 2.31×10^-11^ (Bonferroni correction of 0.001 for 43,222,677 possible interactions) and *HindIII* fragment-level anchors that intersected open chromatin, active histone marks, or brain-expressed TSS consistent with enhancer-promoter (E-P) or promoter-promoter (P-P) interactions (***Methods***). We identified 75,531 HCRCI in adult cortex and 75,246 in fetal cortex (32,007 HCRCI were present at both developmental stages). ***Figure S1*** shows a circular genome plot of these eHi-C readouts from adult and fetal cortex.

**Figure 1.**
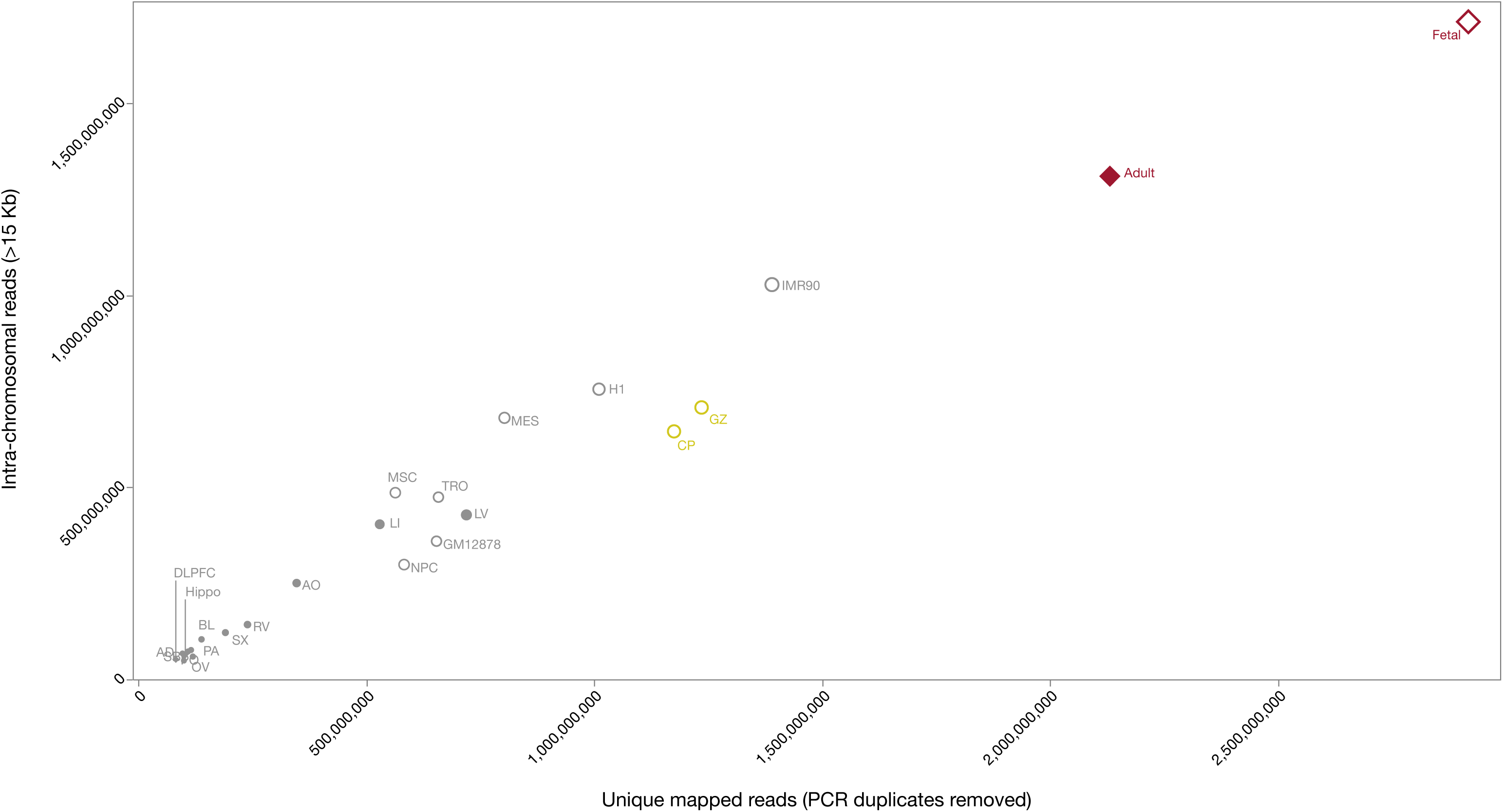
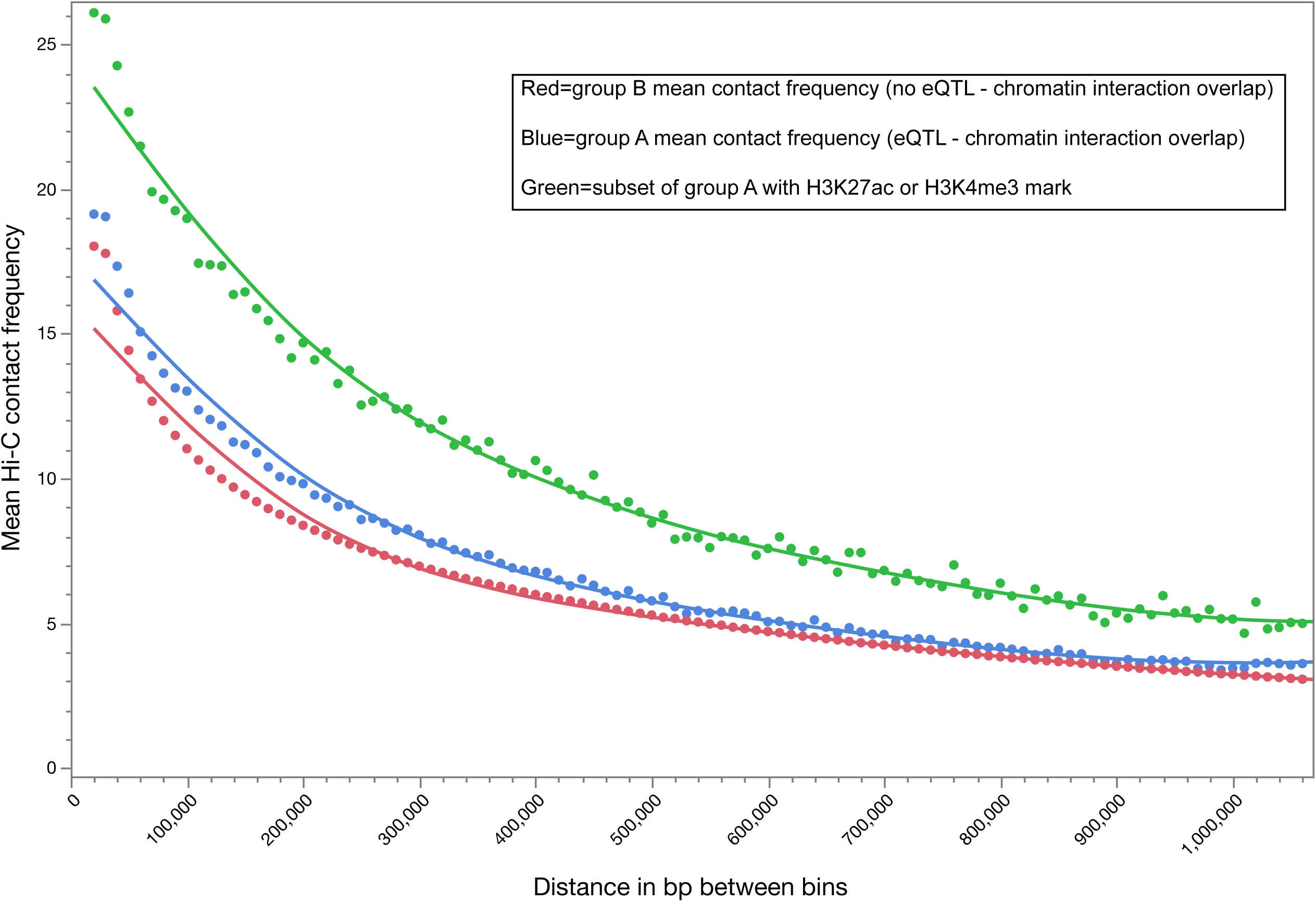
Comparisons of high-level Hi-C metrics and features. <u>Figure 1a:</u> Metrics for 25 Hi-C datasets. The X-axis is the total number of reads passing quality control (uniquely mapped, PCR duplicates removed), and the Y-axis is the number of informative *cis*-reads (uniquely mapped, PCR duplicates removed, intra-chromosomal, >15kb apart). Point sizes are proportional to the numbers of informative *cis*-reads. Red diamonds show data we generated in human brain using eHi-C: filled red diamond is adult temporal cortex (Adult, N=3), and open red diamond is fetal cortex (Fetal, N=3). Yellow open circles show fetal germinal zone (GZ, N=3) and cortical plate (CP, N=3) ^17^. From Schmitt et al. ^32^, filled grey circles show 14 human tissues and 7 cell lines. Tissues: AD=adrenal gland; AO=aorta; BL=bladder; DLPFC=brain dorsolateral prefrontal cortex; Hippo=brain hippocampus; LG=lung; LI=liver; LV=heart left ventricle; OV=ovary; PA=pancreas; PO=psoas skeletal muscle; RV=heart right ventricle; SB=small intestine; SX=spleen. Cell lines: GM12878=lymphoblast; H1=human embryonic stem cell (hESC); IMR90=lung fibroblast; MES=mesoderm; MSC=mesenchymal stem cell; NPC=neural progenitor cell; TRO=trophoblast-like cell. <u>Figure 1b:</u> Brain eQTLs and chromatin interactions. Scatter plot of genomic distance between chromatin interaction anchors (X-axis) and mean Hi-C contact frequency (Y-axis). Using the CommonMind DLPFC eQTL dataset, we stratified by these data when the chromatin interaction anchors did not overlap an eQTL (red), when they did overlap an eQTL (blue). The subset of the chromatin interaction-eQTL overlapping which have H3K27ac or H3K4me3 mark are in green.

We analyzed the properties of these brain eHi-C data to establish a foundation in support of our goal of understanding GWAS results for schizophrenia, intelligence, and other brain traits. These analyses are detailed in the ***Supplemental Note*** and summarized here. First, we compared our eHi-C readouts to external Hi-C datasets for A/B compartments, TAD boundaries, FIREs, and chromatin interactions, and found good agreement with external Hi-C data (***Figures S2-S5***). Second, we evaluated whether these Hi-C readouts captured biologically relevant information (***Tables 1, S3***). We found that FIREs and super FIREs recapitulated key functions of the source tissues: differentiation and neurogenesis in fetal cortex and core neuronal functions in adult cortex. As a control, FIREs and super FIREs from heart ventricle were consistent with basic myocardial functions. GREAT ^45^ analyses of fetal chromatin interactions were enriched for transcriptional regulation and core functions of the major cell types (glia and neurons), and the adult results pointed at postsynaptic density and excitatory synapse. TAD boundaries showed no enrichment consistent with their cell type-independent insulative and structural roles. Third, evaluation of functional genomic features (***Tables 1, S4***) showed that adult FIREs were enriched in adult cortical H3K27ac marks, enhancers, and open chromatin while depleted in H3K4me3 marks ^32^. Fetal brain FIREs were enriched for fetal H3K27ac and CTCF marks. Adult cortex chromatin interactions were enriched for open chromatin, enhancer, CTCF, H3K27ac, and protein-coding TSS while depleted in H3K4me3 marks. In fetal cortex, chromatin interactions were enriched for H3K27ac and CTCF with depletion of H3K4me3 and gene expression. TAD boundaries in adult and fetal brain were enriched for CTCF and TSS of protein coding genes ^34^. Fourth, we evaluated the relation between LD and TADs as both are block-like and span 10^5^-10^6^ bases. TADs are defined by 3D chromatin interactions in cell nuclei whereas LD reflects historical events for a given human sample. We confirmed that TADs and LD blocks identify different genomic regions ^17,46^. Fifth, we conducted evolutionary analyses for brain FIREs and found that these regions have stronger evidence for ancient and recent positive selection, less population differentiation, and fewer singleton/doubleton single nucleotide variants (***Table S5***). These observations suggest that brain FIREs are important genomic regions under stronger population genetic constraints. Sixth, we evaluated the importance of chromatin interactome on gene expression in fetal and adult brain and found: (a) developmentally specific adult vs fetal FIREs had a strong relation to gene expression; (b) in a multivariable model, we found that A/B compartments, FIREs, and chromatin interactions were significant and orthogonal predictors of gene expression (model *R*^*^2^*^ 0.0475, ***Figure S6***); and (c) we found a strong overlap of adult cortex chromatin interactions with adult cortex eQTLs ^18^, particularly for HCRCI (***Figure 1b***).

**Table 1.**
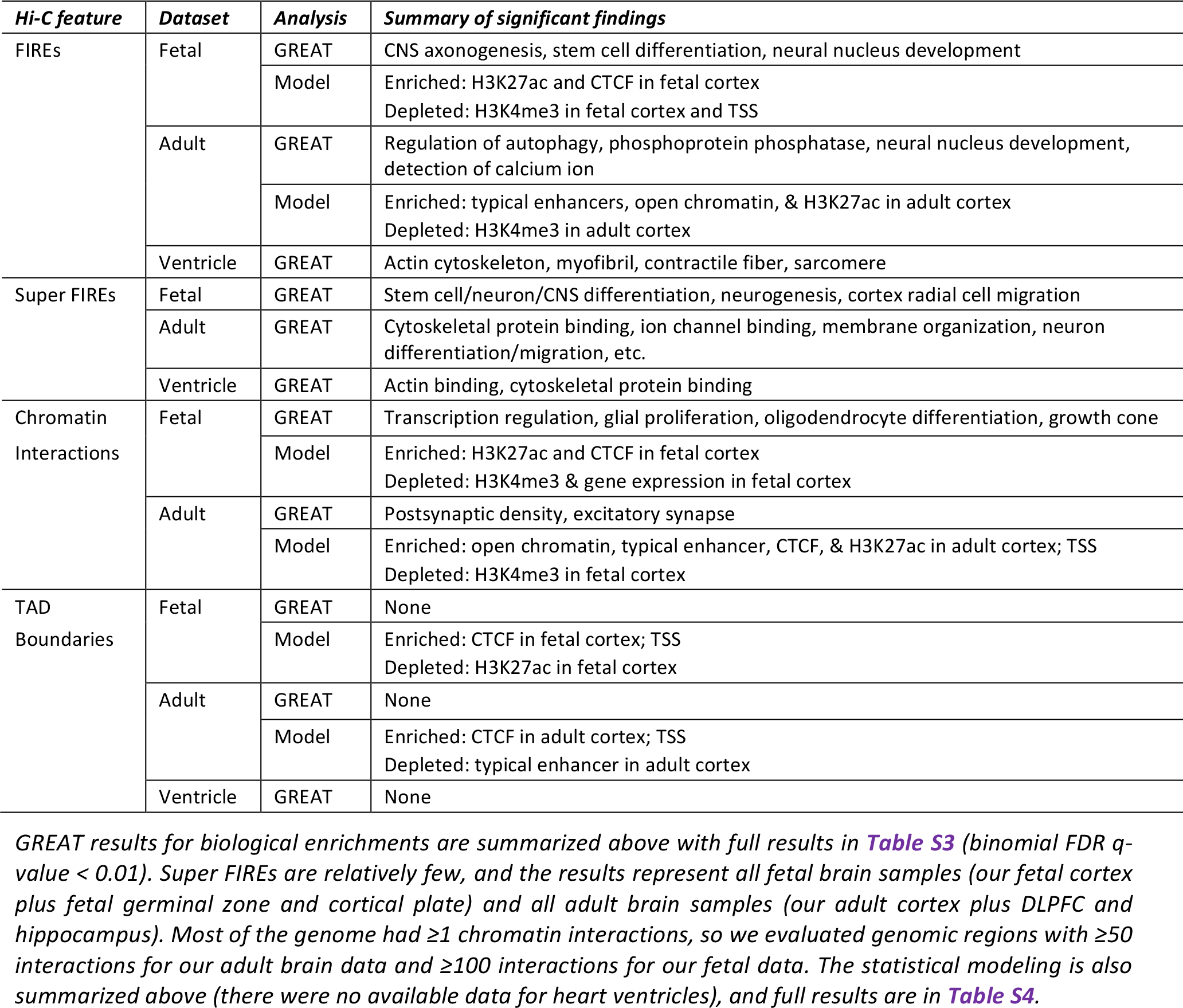
Analysis of Hi-C features

In conclusion, the brain eHi-C data we generated were consistent with prior Hi-C datasets. Although Hi-C data directly incorporate few genomic annotations, Hi-C readouts are an informative functional datatype often orthogonal to other datatypes but with a strong relation to brain gene expression. Evaluating the relation of eHi-C readouts to genetic risk for psychiatric disorders and brain traits is thus empirically-grounded.

### Human GWAS and functional genomics

Having shown the salience of our brain eHi-C data, we now focus on human complex traits. Here, we evaluate GWAS results with respect to tissue-specific gene expression, Hi-C readouts, and brain functional genomic annotations using partitioned LD score regression (pLDSC) ^47^. pLDSC leverages LD, and its use here is appropriate as GWAS results and functional genomic annotations are both at a genome-wide, gigabase scale (LD-based approaches become problematic for individual GWAS loci of 100s of Kb, see below). Although our main interests are schizophrenia and intelligence, we obtained summary statistics for 27 human traits that had been the subject of large GWAS (***Table S2***): 6 psychiatric disorders, 5 CNS traits, 6 neurological conditions, and 10 traits not generally thought to be rooted in the nervous system (e.g., coronary artery disease, total cholesterol).

First, we used GTEx ^43^ bulk tissue gene expression data from 48 regions to identify the tissues implied by the GWAS results (***Figure 2a, Table S6***). These analyses in essence looked for enrichment of SNP-heritability in genes whose expression was specific for a tissue. Of the 27 GWAS, 15 had a significant finding, and the 12 GWAS without a significant finding had relatively few GWAS hits. The control conditions largely corresponded to expectations: coronary artery disease with aorta, tibial, and coronary artery; hemoglobin A1c with whole blood; inflammatory bowel disease with whole blood and spleen; total cholesterol with liver; waist:hip ratio with adipose; age at menarche with uterus; and body mass index with brain ^48^. Coronary artery disease, height, and inflammatory bowel disease had other significant enrichments for tissues not immediately known to be involved – we believe this resulted from cell type mixtures within the bulk tissues studied. The psychiatric disorders/CNS traits with the largest number of GWAS hits (schizophrenia, major depression, bipolar disorder, intelligence, educational attainment, and neuroticism) enriched for multiple brain regions with the strongest associations usually for brain cortex, frontal cortex, and nucleus accumbens. Gene expression specificity findings are consistent with the high genetic correlations between these traits ^16^. None of the neurological conditions had a significant association, likely due to the small sizes of the primary studies.

**Figure 2.**
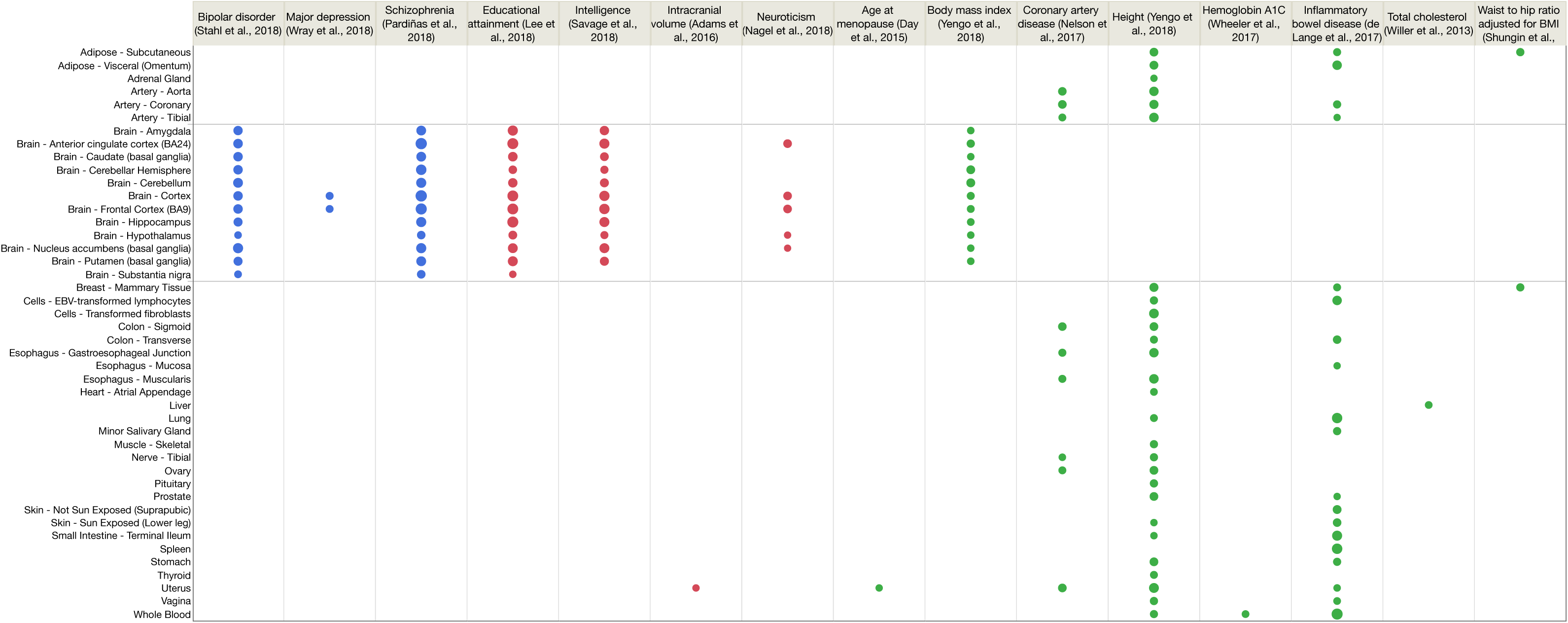
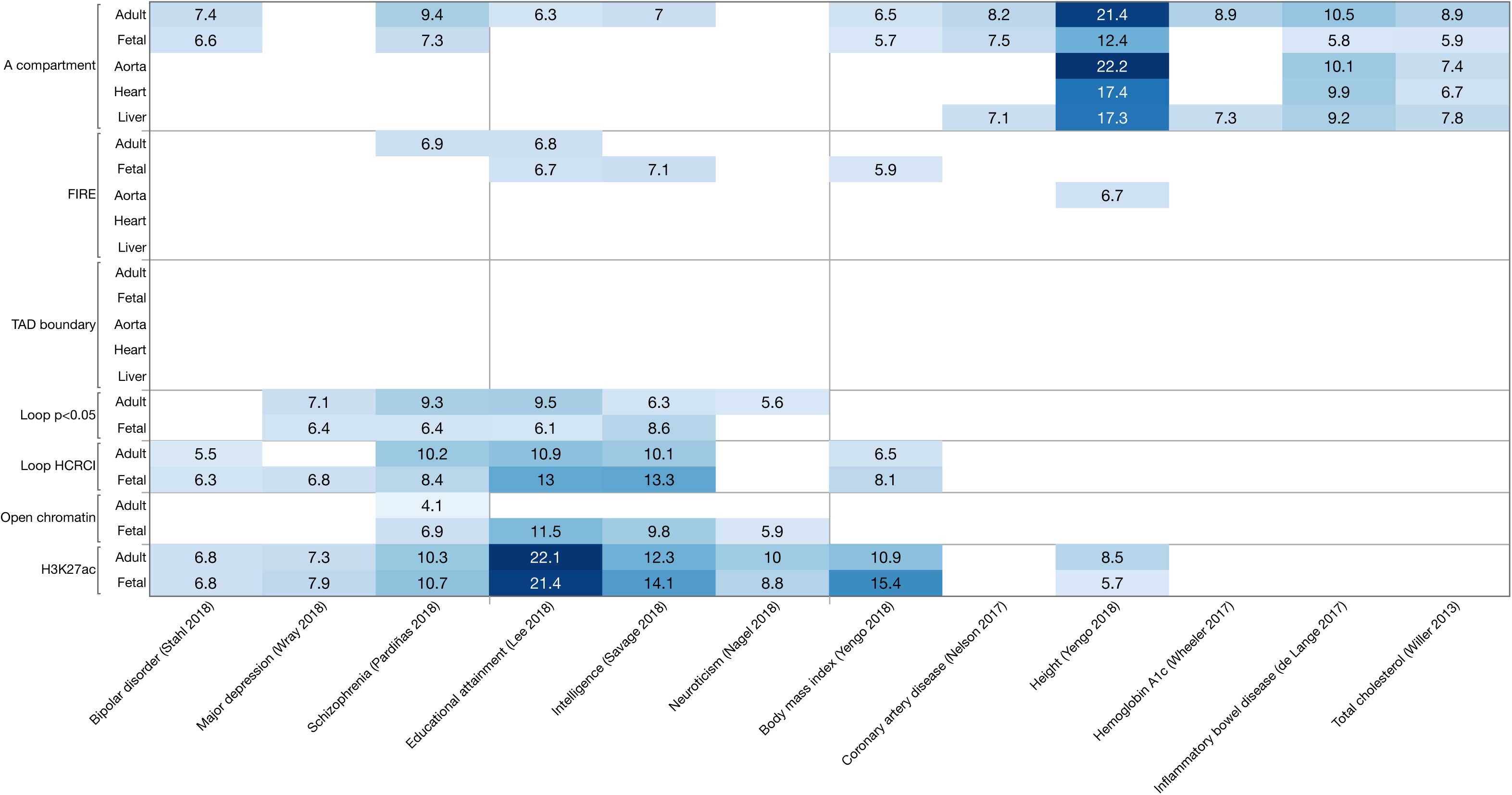
Connecting GWAS summary statistics to tissues using pLDSC and bulk RNA-seq data from GTEx. <u>Figure 2a:</u> Results of analyses connecting GWAS summary statistics to tissues using bulk-tissue RNA-seq using pLDSC. RNA-seq data were from 48 GTEx tissues (v7, after removing 5 tissues with N<100). The significance level was 0.001 corrected for 1296 comparisons (27 GWAS x 48 tissues), or *P* < 7.72e-7. The panels along the X-axis show the GWAS trait. The Y-axis shows GTEx tissues. The circles show significant trait-tissue associations. Circle sizes are proportional to –log10(P). Blue circles are psychiatric disorders, red circles are for brain trait, and green circles are other traits and diseases. For clarity, traits with no significant associations are omitted from the figure (ADHD, age at menarche, Alzheimer’s disease, amyotrophic lateral sclerosis, anorexia nervosa, autism spectrum disorder, epilepsy, hippocampal volume, Parkinson’s disease, stroke, and type 2 diabetes). <u>Figure 2b</u> depicts a heat map for results of partitioned LD score regression for seven large GWAS for functional genomic readouts (Hi-C, ChIP-seq, gene expression, and open chromatin) for multiple tissues. See Figure 1a legend for abbreviations.

Second, we evaluated SNP-heritability enrichments of these GWAS with functional genomic data including Hi-C, open chromatin, and H3K27ac histone marks (***Figure 2b*** and ***Table S7***). Chromatin interactions had notable SNP-heritability enrichment for psychiatric disorders and CNS traits – particularly the anchors of HCRCI (adjusted P<0.001 and an E-P or P-P chromatin interaction) which were a subset of all significant chromatin interactions (adjusted P<0.05). H3K27ac marks showed enrichments for most brain traits but also for BMI and height. There was a tendency for stronger SNP-heritability enrichment in functional annotations in adult brain for schizophrenia, and in fetal brain for educational attainment and intelligence. FIREs had modest enrichments without clear specificity by organ or developmental stage. In contrast, TAD boundaries showed no significant SNP-heritability enrichments and open chromatin compartments (“A”) had significant enrichments for many traits and tissues.

From these results, we conclude that using functional genomic data from adult cortex is a reasonable choice for evaluation of schizophrenia and intelligence, particularly when complemented by fetal cortical data. The functional annotations that showed the most specific connections to schizophrenia and brain traits were open chromatin and HCRCI. Thus, genome-wide annotations that capture dynamic genome processes central to gene regulation in brain have particular salience to genetic risk for multiple brain-based traits.

### Interpreting GWAS loci using a 3D perspective on brain nuclear architecture

The preceding sections support our choices of tissue (adult and fetal cortex) and salience of functional genomic readouts for interpretation of GWAS findings for schizophrenia, intelligence, and other brain traits. In this section, we attempt to “connect” significant GWAS loci to specific genes as this is a crucial deliverable of GWAS for idiopathic disorders of the brain. Our approach had the following steps: (a) curate all genome-wide significant loci for schizophrenia, intelligence, and eight other CNS traits; (b) as a special case, identify statistically independent loci functionally connected to the same gene; (c) as a special case, evaluate GWAS loci that intersect no genes (intergenic) or a single gene to test the strong assumption that GWAS findings connect to the nearest or intersecting gene; (d) identify all genes implicated by location, HCRCI, and/or eQTL evidence; and (e) contrast the biological pathways implicated by these methods of assigning associated SNPs to genes.

We focused on cortex given empirical data connecting this tissue to schizophrenia and intelligence using orthogonal functional genomic data (bulk tissue mRNA-seq, single-cell RNA-seq, enhancer marks, and open chromatin) ^28,49^-^51^. We selected an inclusive set of 16,308 genes (77.8% of all protein-coding genes, GENCODE) with any expression in adult or fetal cortex ^43,44^. Second, we selected eQTL SNP-gene pairs from CommonMind or GTEx (*q*<0.05) ^18,43^. Third, using our eHi-C data, we identified HCRCI in adult or fetal cortex (*P*<2.31×10^-11^, Bonferroni correction of α=0.001 for 43,222,677 possible interactions). As in ENCODE and PsychENCODE ^19,52^, we identified anchors that overlapped enhancers (E) or promoters (P) using cortical functional genomic data from the same developmental stage (***Table S2***). E were defined as the intersection of: eHi-C *HindIII* fragment within an anchor, open chromatin, and either a H3K27ac peak or a H3K4me3 peak overlapping the start site of a brain-expressed transcript. P were defined as brain-expressed transcripts overlapping open chromatin. We focused on 75,531 adult and 75,246 fetal cortex E-P or P-P HCRCI. ***Figures 8a-g*** show representative examples as browser tracks.

First, we obtained results for the most recent GWAS of schizophrenia, ADHD, alcohol dependence, Alzheimer’s disease, anorexia nervosa, autism spectrum disorder, bipolar disorder, and major depression along with intelligence and educational attainment (***Table S2***). Loci were defined similarly across studies using LD-based “clumping”, merging of loci ±50 Kb, and removal of loci with a single significant SNP. Each locus is an LD-based genomic region containing multiple genome-wide significant SNP associations for one of these 10 traits. There were 938 loci across these 10 traits: ≤12 for alcohol dependence, autism, anorexia nervosa, and ADHD; 24 for Alzheimer’s disease; 29 for bipolar disorder; 44 for major depression, 145 for schizophrenia; 205 for intelligence; and 467 for educational attainment. The median *P*-value per locus was 1.17e-9 (interquartile range, IQR 1.19e-8 – 1.21e-11). The loci had a median size of 197 Kb (IQR 87-424 Kb), and 21.9% were intergenic, 38.4% intersected a single gene, and 39.8% were multigenic (containing a median of 4 genes, IQR 2-8). As anticipated from the primary studies, overlap was common as 39.4% of the loci were for ≥2 disorders.

Second, we identified genes that were implicated by different statistically independent loci for each trait. (a) As listed in ***Table S8a***, 16 genes had two statistically independent trait associations located >100 Kb apart for a trait (e.g., *RBFOX1* had two hits separated by >1.2 Mb for both major depression and intelligence, and *DPYD*, ***Figure S8a***). (b) We found that 40 eQTL-genes had an eQTL-SNP in statistically independent loci >100 Kb apart (30 genes for educational attainment, 7 for schizophrenia, and 3 for intelligence, ***Table S8b***). For example, the expression of *SATB2* in adult cortex is significantly associated with SNPs in schizophrenia loci located 242 Kb apart (***Figure S8b***). (c) Unexpectedly, brain Hi-C data showed that some LD-defined loci nonetheless contained evidence of functional interactions (***Table S8c***). The presence of many bridging HCRCI suggests that these statistically independent loci have functional regulatory activity in nuclei from adult and fetal cortex (e.g., 9 genes in three schizophrenia loci from chr2:198.1-201.3 Mb may form a functionally connected unit, ***Figure S8c***).

Third, we evaluated 199 intergenic loci and 360 loci that intersected a single gene (excluding the extended MHC region). Intergenic loci were a median of 191.7 Kb from the nearest protein-coding gene (IQR 69.2-412.5 Kb), and total-stranded RNA-seq from an adult and a fetal cortical sample did not reveal unannotated transcripts. The standard assumption is to assign a GWAS locus to the nearest gene: based on brain eQTL and HCRCI, this assumption was strictly correct for only 19 loci (9.6%), partly correct for 25 loci (12.6%, the nearest gene was implicated but so were 1-9 additional genes), incorrect for 47 loci (23.6%, eQTL and/or HCRCI connected to 1-7 genes that were not the closest to the locus), and there were no eQTL or HCRCI connections for 108 loci (54.3%). For single gene loci, the standard assumption is to assign the GWAS locus to the gene it intersects. This assumption was strictly correct for 112 loci (31.1%), partly correct for 208 loci (57.8%, the intersecting gene was implicated but so were 1-37 additional genes), and incorrect for 19 loci (5.3%). Genes implicated via eQTL or HCRCI are often far from a locus and beyond typical LD block sizes: a median of 543.1 Kb (IQR 146.9-896.2 Kb) from intergenic loci and 409.1 Kb (IQR 171.5-825.2 Kb) from single gene loci. Thus, functional genomic data from human brain indicate that typical assumptions used in bioinformatics analyses of GWAS results are usually incorrect or oversimplifying.

Fourth, we systematically evaluated GWAS results using functional genomic data. We used three approaches to create explicit hypotheses for subsequent biological experiments: (a) implication by genomic location, that significant SNPs in a GWAS locus implicate the genes it intersects (directly or by LD-proxy); (b) eQTL pairs consist of an eQTL-SNP in a GWAS locus that is significantly associated with the expression level of an eQTL-gene that may or may not be in the locus; and (c) HCRCI capture the regulatory potential created by physical proximity of two genomic regions in brain nuclei: one anchor is in a GWAS locus and the other anchor is in a gene that may or may not be in the locus (HCRCI have properties consistent with E-P or P-P regulation). The latter two methods are based on functional genomic data from adult and fetal cortex. We did not apply statistical prioritization algorithms (e.g., TWAS, co-localization, or credible SNP) as these were not aligned with our goal of generating an inclusive set of locus-to-gene hypotheses. Some methods are highly restrictive (e.g., only 69 of 18K significant SNPs in CLOZUK were “credible” with posterior probabilities >0.9) and, to our knowledge, these methods have not been rigorously benchmarked in experimental models.

We compared our results to Wang et al. ^19^ on the same schizophrenia results ^12^ (***Methods***). There was substantial overlap with 81.0% (596/736) of the protein-coding genes identified by Wang et al. also found by us, particularly for their “high-confidence” genes (95.9%, 260/271). However, our method identified many more genes (1,300 vs 736) mainly due to our far deeper eHi-C data.

Encouraged by this overlap, we applied our strategy to 938 significant GWAS loci for 10 psychiatric disorders and brain traits. Using location, eQTL, and/or HCRCI, we connected these loci to 8,595 genes. The implicated genes are listed in ***Table S9***. These genes had the expected locus type distributions: 84.7% in multigenic loci, 13.0% in single gene loci, and 2.3% in intergenic loci. ***Figure 3a*** shows the number of genes identified per trait (range 12-3,657) which, as expected, was highly correlated with the number of significant loci (Spearman ρ=0.988). Many genes were identified by ≥2 traits (42.1%, ***Figure 3b***) consistent with the primary papers that reported considerable locus overlap for many traits. ***Figure 3c*** is a Venn diagram of the interrelations of the three methods of connecting GWAS loci to genes. Genes identified by location usually (94.4%, 3,091/3,276) also have eQTL (58.1%) and/or HCRCI evidence (86.9%). However, location provides a simplified portrait of the genes involved with respect to functional genomic data from human brain: 3,276 genes are implicated by location (38.1%), 3,121 genes by eQTLs (36.3%), and 7,581 by HCRCI (88.2%). See ***Figures S8d-g*** for examples. Genomic location appears to be a specific but insensitive indication of the involvement of a gene.

**Figure 3.**
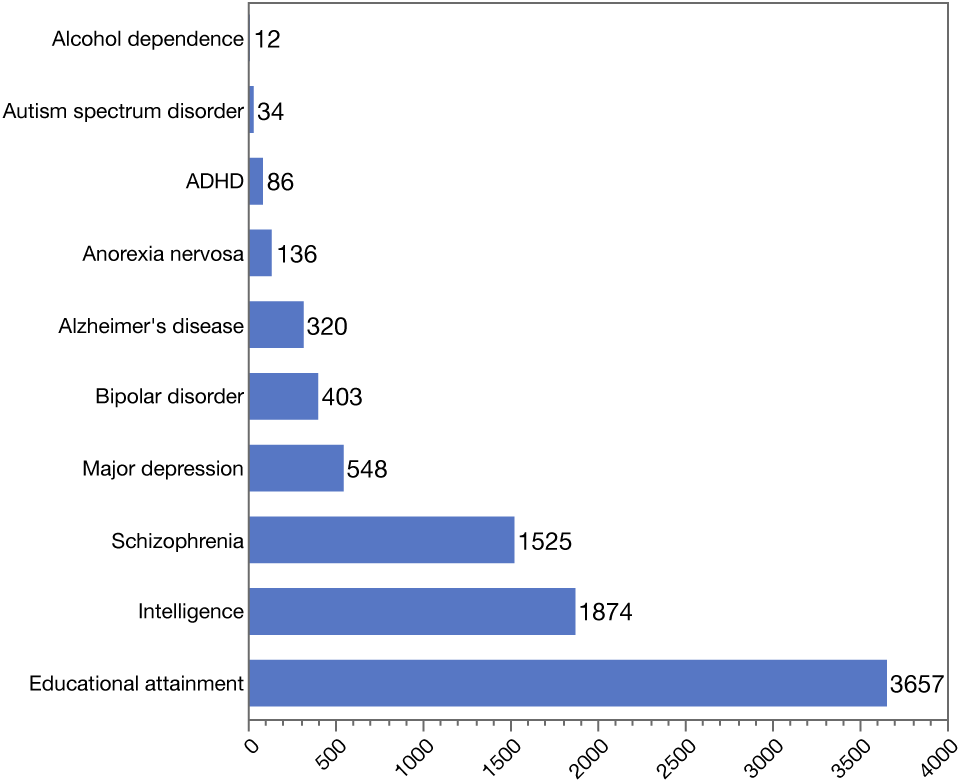
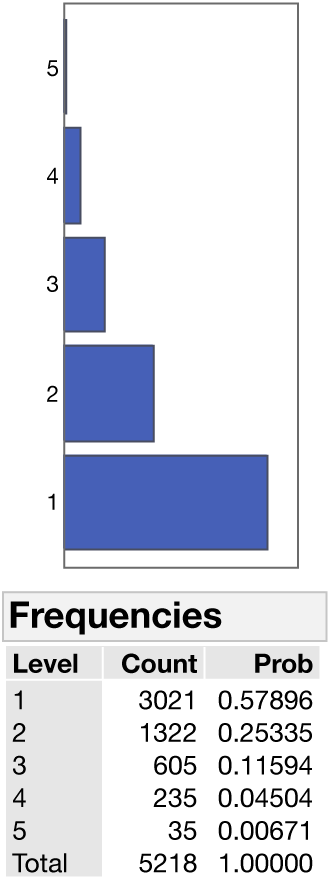
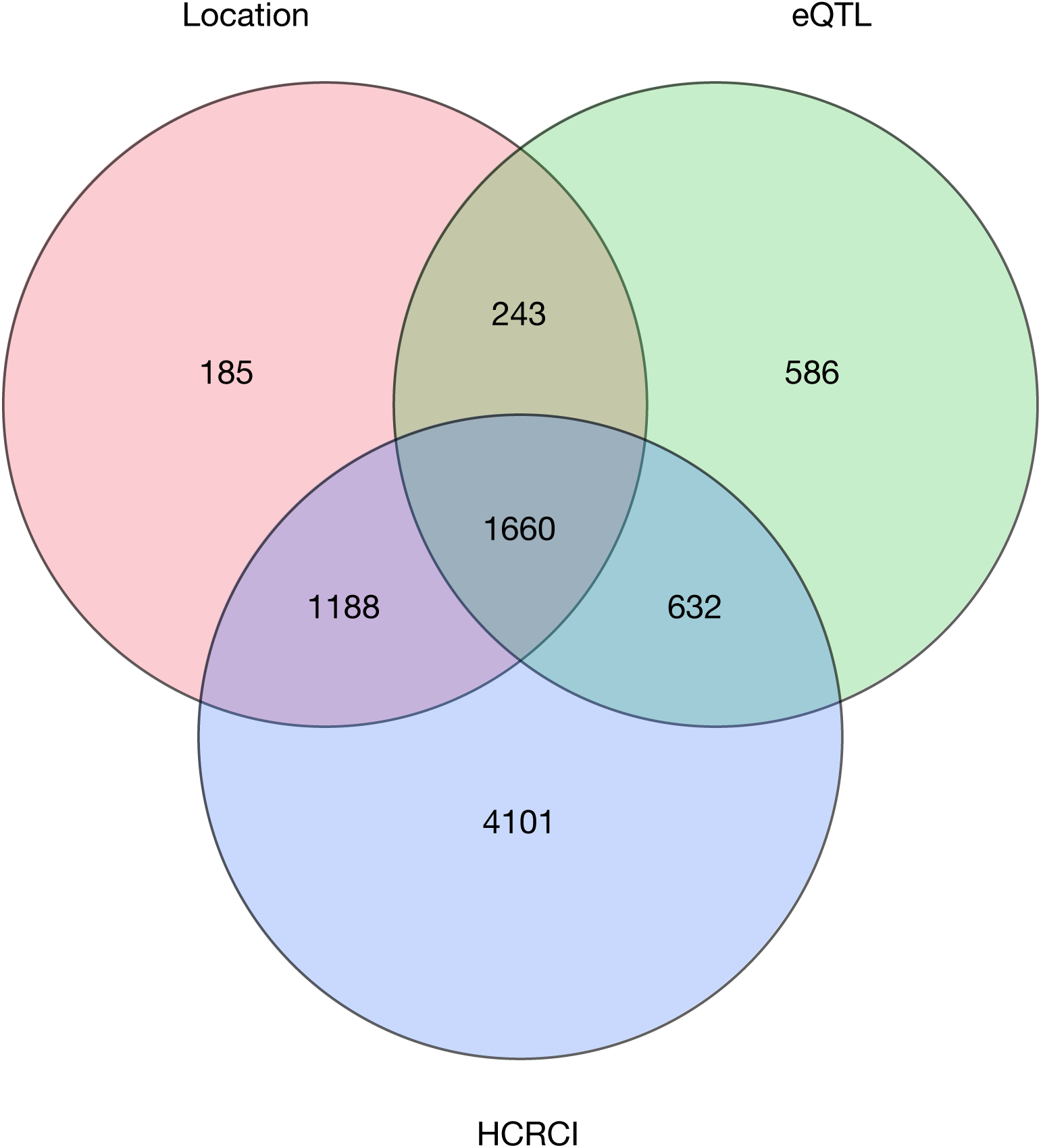
HCRCIs provide the most comprehensive viewpoint of SNP-to-gene connections. <u>Figure 3a:</u> Histogram of the number of genes identified using location, eQLT, or HCRCI evidence for each of the 10 brain disorders, diseases, or traits. <u>Figure 3b:</u> Histogram and percentages for the number of times a gene was identified. A total of 5,218 genes were implicated, 57.9% genes for one of the ten traits and 42.1% for two or more traits. <u>Figure 3c:</u> Venn diagram of the number of genes implicated by location in a GWAS locus (Location), by brain cortex eQTL data, or by HCRCI. The latter dominates. Genes implicated by location usually have eQTL or HCRCI evidence as well but sole reliance on location yields an overly simplified view of the potential complexity of the implicated biology.

Fifth, for schizophrenia and intelligence, given clear evidence of polygenicity (that “many genes” are involved), a crucial unanswered question is enumeration of the biological processes implied by the identified sets of genes. Identification of biological processes has major implications for development of unifying etiological hypotheses and therapeutic strategies. Many statistical approaches to gene set analysis are based on proximity and/or LD ^53^, but functional genomic readouts (eQTL and HCRCI) are based on regulatory processes. We conducted a comprehensive set of gene set analyses using hypergeometric rather than LD-based tests. The gene sets evaluated are listed in ***Table S2***, and include standard gene sets as well as gene sets previously found to be important or of interest to us ^10,54,55^. Location, eQTL, or HCRCI implicated 45 gene sets for schizophrenia (***Table S10***) and 46 gene sets for intelligence (***Table S11***). For both schizophrenia and intelligence, most gene sets were implicated either by location or HCRCI alone. Crucially, biological inference differed: location evidence suggested gene sets relating to transcript binding and neuronal/synaptic processes whereas HCRCI implicated more fundamental regulatory/chromatin biology. For intelligence, there were exceptionally strong connections to two overlapping cell adhesion gene sets (*P*<1e-300), and the 169 brain-expressed genes included neurexin 1, neuroligin 1, multiple cadherin genes, and genes relevant to developmental neuronal migration; these gene sets connect genetic variation in human intelligence to neuronal migration and establishment of synaptic contacts.

Finally, 3 genes have notable connections to specific brain maladies: Alzheimer’s disease-*APOE*, alcohol dependence-*ADH1B*, and schizophrenia-*C4A*. Each of these genes is in the most significant GWAS locus for that disorder, and additional data implicate the protein product. Integrating functional genomic data, however, suggests that a sole focus on these single genes may be oversimplifying – e.g., eQTL or HCRCI implicate many genes outside the GWAS locus for Alzheimer’s disease (37 genes) and alcohol dependence (5 genes), and *C4A* has a dense pattern of HCRCI to other genes. Therefore, although the pathological insights provided by these single genes may be important, an exclusive focus on these genes may be inadequate to fully understanding the etiological processes implied by the GWAS findings.

### A taxonomy of TADs

In studying the schizophrenia results in ***Table S10***, we observed that genes implicated by HCRCI had significantly greater expression in adult cortex (*R*^*^2^*^=0.0058, *P*=1.64e-21) compared to genes identified by location (*R*^*^2^*^=0.0014, *P*=1.45e-6) or eQTLs (*R*^*^2^*^=0.0013, *P*=7.84e-6). Inspection of these results in genomic context suggested a relation to adult brain TADs as 11.9% of these TADs contained an association with schizophrenia. We note that most CLOZUK loci were entirely within an adult brain TAD (136/145, and 8 loci spanned 2 TADs and the MHC locus encompassed 3 TADs). It was readily apparent that these TADs had markedly different properties leading us to investigate quantitative characterization of TADs. As described in the ***Supplemental Note***, we applied PCA to six features for each adult TAD (gene density plus measures of adult cortical function including gene expression and four adult brain Hi-C readouts). None of these six features directly index schizophrenia risk. PC1 captured 44.9% of the variance, and the item loadings indicated that PC1 reflected active functional status.

PC1 was greater in TADs with versus without a significant schizophrenia association (*P*=0.008). PC1 had modest predictive capacity for whether genetic variants within a TAD would become genome-wide significant in subsequently larger studies (using 3 schizophrenia GWAS: PGC1 9,394 cases, 7 loci; PGC2 36,989 cases, 108 loci; and CLOZUK 40,675 cases, 145 loci) ^11,12,56^. TADs that did not contain a significant association in PGC1 but which subsequently became significant had greater mean PC1: *P*=0.047 comparing PGC1 to PGC2, and *P*=0.0095 for PGC1 to CLOZUK.

We converted PC1 to deciles. Of the 105 TADs in the top decile for PC1, 22.9% contained a genome-wide significant association for schizophrenia. The highest PC1 decile was significant enriched for schizophrenia SNP-heritability (pLDSC enrichment 1.64, SE=0.113, *P*=5.53e-8), the lowest decile was significantly depleted for schizophrenia SNP-heritability (pLDSC enrichment 0.68, SE=0.062, *P*=9.19e-7), and all other deciles showed no significant enrichment (***Figure S7***). Intriguingly, we observed a similar pattern of SNP-heritability enrichment in the highest decile with depletion in the lowest decile for diverse human traits (body mass index, total cholesterol, height, hemoglobin A1C, and inflammatory bowel disease) even though PC1 was mainly based on brain functional genomic data. TADs in the top decile were broadly expressed in adult organs (mean of 21.0 of 30 GTEx organs) compared to TADs in the bottom decile (mean 5.4). TADs in the top decile were more likely to have significant associations – excluding psychiatric disorders – in the NHGRI-EBI GWAS catalog (mean of 13.8 associations vs 1.8 in the bottom decile). The proportions of genes in a TAD that are intolerant to loss-of-function variation (pLI>0.9) ^57^ and which were specific to pyramidal neurons and medium spiny neurons (the brain cell types we previously identified to be enriched for schizophrenia heritability) ^49^ were greatest in the second and third deciles and lower in the top or bottom deciles.

As an example, ***Figure S9*** shows the TAD with this highest PC1 value (chr11:63.98-67.38 Mb). This region contains two significant loci for bipolar disorder and a significant locus for schizophrenia. Most of the TAD is a FIRE or super FIRE, and it is dense with adult cortex eQTLs and HCRCI. The TAD contains dozens of genes expressed at high levels in many tissues. It contains many significant associations from GWAS as well as multiple disease connections in OMIM.

Thus, TADs are heterogeneous with respect to functional characteristics and disease salience. TADs with the most genes and greatest functional indices tended to contain genes expressed in most of the body along with GWAS associations for many different traits. Top decile TADs are large, complex multigenic regions. However, the lower end of the functional spectrum tended to have intergenic and single-gene associations that may provide easier entrée into molecular networks underlying schizophrenia.

### Rare genetic variation associated with neuropsychiatric disorders

#### Exonic variation

We evaluated the salience of Hi-C readouts for genes implicated in rare variant studies of intellectual disability (i.e., the lower tail of the cognitive ability distribution). We compiled a gene set from literature reviews, OMIM, and exome sequencing studies ^58^-^60^. We could not analyze schizophrenia because too few genes have been identified. Full details of this analysis are in ***Table S12***. There were significant univariate associations between the implication of a gene in intellectual disability via exonic variation and its intersection with FIREs, TAD boundaries, or the number of chromatin interactions in both adult and fetal eHi-C datasets. In a multivariable model for the adult eHi-C data, intersection with a TAD boundary increased the odds of being an intellectual disability gene by 54% (OR=1.54, *P*=2.56e-4) as did the number of chromatin interactions (OR=1.26 for each doubling of the number of interactions, *P*=6.61e-11). Intersection with an adult FIRE was not significant in this model. In a multivariable model for the fetal eHi-C data, the odds of being an intellectual disability gene increased markedly with the number of chromatin interactions (OR=4.11 for each doubling of the number of interactions, *P*=8.0e-13) but fetal FIRE and TAD boundary intersections were not significant in this model.

#### Copy number variation (CNVs)

Cases with schizophrenia have more rare CNVs than controls ^61^. Excess CNV “burden” in cases can be attributed to genic CNVs with greater effects for deletion than duplication CNVs ^62,63^. Greater CNV burden in cases persists after removing ~10 large CNVs individually associated with schizophrenia (e.g., 22q11 or 16p11) ^62^. It is plausible that a CNV that disrupts a FIRE or a TAD ^64^ could explain some of the excess CNV burden. We evaluated this hypothesis using carefully curated CNVs in 4,719 schizophrenia cases and 5,917 controls ^63,65^. We excluded large CNVs known to be associated with schizophrenia ^62^, and controlled for genotyping batch and ancestry. The presence of a CNV deletion intersecting one or more adult brain FIREs was significantly associated with schizophrenia (OR=1.72, P=3.27e-6) whereas CNV duplications were not associated (OR=1.10, *P*=0.17). The presence of a CNV deletion intersecting one or more fetal brain FIREs was modestly associated with schizophrenia (OR=1.31, *P*=4.7e-3) as were CNV duplications (OR=1.22, *P*=7.9e-3). CNVs that intersected TADs were not notably associated with schizophrenia: adult TAD-CNV deletion OR=1.08 (P=0.60); adult-TAD CNV duplication OR=1.16 (*P*=0.047); fetal TAD-CNV deletion OR=1.07 (*P*=0.64); and fetal-TAD CNV duplication OR=1.16 (*P*=0.042).

## Discussion

Our understanding of human genetic variation is currently best at the extremes. At a chromosomal scale, there is considerable knowledge of the prevalence and medical relevance of variation (i.e., large structural variants). At the base pair scale, we have increasingly detailed surveys of the nature and frequency of genetic variation from studies like TOPMed and gnomAD. Between these extremes, some annotations appear to be increasingly complete for a few crucial topics (e.g., gene models, variation causal for single-gene diseases, or expression patterns in human tissues). However, particularly for complex psychiatric disorders of profound societal importance, there is an unsolved problem at intermediate scales: given the typical paucity of exonic findings, precisely how do the thousands of significant, subtle, and common associations that account for most of inherited liability act mechanistically to increase risk for disease?

In this paper, we used deep eHi-C datasets from human cortex to evaluate its utility to systematically connect GWAS loci to genes. This is a key deliverable of these genetic studies, to provide an enumeration of the genes implicated which are difficult to obtain otherwise. Our approach was somewhat different from prior efforts in that we carefully evaluated our data and their salience prior to the major component of the paper, systematic evaluation of GWAS for seven psychiatric disorders, Alzheimer’s disease, intelligence, and educational attainment. Instead of only considering the genome as a 1D object defined by LD relationships, we used a 3D functional snapshot of genome organization in brain cells. Moreover, we sought to delimit complexity via a fuller evaluation of all implicated regions (e.g., not focusing on a far smaller set of “credible” SNPs). In doing so, we sought to identify a more complete parts list. We used this strategy to connect 938 significant GWAS loci for 10 brain traits to 8,595 genes. These connections provide solid hypotheses for subsequent biological experiments.

We found that genomic location is a problematic way to understand the genes implicated by GWAS ^19^. LD is a fundamental feature of the genome with a large body of supporting statistical genetic theory and analytical methods ^66,67^. Indeed, LD has been essential to genetic discovery for decades: crucial to linkage mapping for Mendelian diseases in large affected pedigrees and a fundamental reason why GWAS “works” (the information in millions of polymorphic SNPs can be captured by many fewer SNPs). LD is a double-edged sword: following identification of a significant locus for a complex disease, LD almost always confounds attempts to identify specific genes. There usually are many loci with approximately the same significance values and, even with dense additional genotyping, rarely can a single variant with markedly greater significance be identified. Leveraging 1D epigenetic information can help in prioritization ^52^ although many epigenetic marks are common (e.g., brain open chromatin regions are about as large as the exome) ^50^, and LD remains a significant hindrance with large number of significant SNPs in high LD that overlap promoters, open chromatin, and histone marks. Difficulties of LD-based approaches following locus identification in complex disease warrant incorporation of more informative data types. LD arises from historical population genetic processes whereas the chromatin interactome captures the functional organization of cells in a disease-relevant tissue. These two features do not overlap well. Our results provide support for the idea that, following genetic identification of a locus for a complex disease like schizophrenia, it is essential to incorporate knowledge of the 3D interactome in a disease-relevant tissue. Connecting GWAS findings to genes using 1D-based methods like LD is often misleading given that the genes implicated by 3D interactome are not in LD or not the nearest gene.

Above and beyond efforts to connect loci to genes, we identified a widespread phenomenon – there exist dozens of genomic regions within functionally-defined TADs that contain large numbers of genes expressed in brain and enriched for schizophrenia associations. Surprisingly, these regions were also enriched for many other brain and non-brain diseases and contained many genes with broad expression patterns. Within these high-activity megabase-scale TADs, however, different genes were implicated for different traits. We suggest that these observations expose a phenomenon for which we currently have no theory, the molecular mechanisms by which GWAS findings for complex disorders alter the coordinate regulation of genes in these dense, highly active genomic regions. This observation has a superficial similarity to the “omnigenic” model ^68^ (but see also ^69^), but differs as we explicitly suggest that GWAS signal is concentrated in identifiable high-activity TADs that are common across multiple disorders as well as in different TADs of simpler and more accessible functional architectures. We do not provide a solution to this complex topic; however, we suggest that articulating the questions and acknowledging the complexity is a key initial step toward deriving a solution.

## Online Methods

### General & data availability

All procedures on data from human research subjects were approved by the appropriate ethical committees. All genomic coordinates are given in NCBI Build 37/UCSC hg19.

Upon acceptance of this paper, eHi-C readouts will be posted on the PGC website and GEO and made available in FUMA and HUGIn (***URLs***).

### Samples

Anterior temporal cortex was dissected from postmortem samples from three adults of European ancestry with no known psychiatric or neurological disorder (Dr Craig Stockmeier, University of Mississippi Medical Center). Cortical samples from three fetal brains were obtained from the NIH NeuroBiobank (gestational age 17-19 weeks), and none were known to have anatomical or genomic disease. Samples were dry homogenized to a fine powder using a liquid nitrogen-cooled mortar and pestle. All samples were free from large structural variants (>100 Kb) detectable using Illumina OmniExpress arrays. Genotypic sex matched phenotypic sex for all samples.

### Easy Hi-C (eHi-C) methods

We used eHi-C to assess chromatin interactome ^31^. The eHi-C protocol is biotin-free and uses sequential enzymatic reactions to maximize the recovery of DNA products from proximity ligation. The main advantage of this Hi-C adaptation is that it can generate Hi-C libraries that are comparable to traditional Hi-C but with lower sample input (as little as 10 mg brain tissue) and increased yield. All of these features of eHi-C are crucial for relatively uncommon human postmortem brain samples.

We followed the protocol described in Lu et al. ^31^. Pulverized tissue (~100 mg) was crosslinked with formaldehyde (1% final concentration) and the reaction was quenched using glycine (150 mM). We lysed samples on ice with brain tissue-specific lysis buffer (10 mM HEPES; pH 7.5, 10 mM KCl, 0.1 mM EDTA, 1 mM dithiothreitol, 0.5% Nonidet-40 and protease inhibitor cocktail), Dounce homogenized, and digested using the six base pair restriction enzyme *HindIII*. This was followed by *in situ* ligation. Samples were reverse cross-linked with proteinase K and purified using phenol-chloroform. DNA was then digested with four base pair restriction enzyme *DpnII* followed by size selection using PCRClean DX beads (Aline Biosciences) (choosing fragments between 100-1000 bp). The DNA products were self-ligated overnight at 16°C using T4 DNA ligase. Self-ligated DNA was purified with phenol-chloroform, digested with lambda exonuclease, and purified using PCRClean DX beads. For re-linearization of circular DNA, bead-bound DNA was eluted and digested with *HindIII* and purified using PCRClean. Bead-bound DNA was eluted in 50 μl nuclease-free water. Re-linearized DNA (~50 ng for one library) was used for library generation (Illumina TruSeq protocol). DNA was end-repaired using End-it kit (Epicentre), “A-tailed” with Klenow fragment (3′–5′ exo–; NEB), and purified with PCRClean DX beads. The 4 μl DNA product was mixed with 5 μl of 2X quick ligase buffer, 1 μl of 1:10 diluted annealed adapter and 0.5 μl of Quick DNA T4 ligase (NEB). Ligation was done by incubating at room temperature for 15 minutes. DNA was purified using DX beads, and eluted using 14 μl nuclease-free water. To sequence eHi-C libraries, we used custom TruSeq adapters in which the index is replaced by 6 base random sequences. Libraries were then PCR-amplified and deeply sequenced (2-5 independent libraries/sample) using Illumina HiSeq4000 (50 bp paired-end).

Because nearly all mappable reads start with the *HindIII* sequence AGCTT, we trimmed the first 5 bases from every read and added the 6-base sequence AAGCTT to the 5’ of all reads. These reads were aligned to the human reference genome (hg19) using Bowtie ^70^. After mapping, we kept reads where both ends were exactly at *HindIII* cutting sites. PCR duplicates with the same positions and UMI were removed ^31^. We also removed read pairs with the two ends within the same *HindIII* fragment. To further filter valid ligation products from *cis*-*HindIII* pairs, we split reads into three classes based on strand orientation: “same-strand” had both ends on the same strand; “inward” had the upstream end on forward strand; and “outward” where the upstream end was on the reverse strand ^31^. “Outward” read pairs with gap distance <1 Kb between the two corresponding fragments were removed because they might originate from undigested *HindIII* sites. “Inward” read pairs with gap distance <25 Kb between the two corresponding fragments were removed because they might come from self-circled DNA with undigested *HindIII* sites. All other reads are valid ligation products and were processed as described below (FIREs, chromatin interactions, TADs, and compartment A/B designation).

### Hi-C readouts

We adapted in-house pipelines to process eHi-C and conventional Hi-C data from external datasets as described previously ^32,71^ with slight modifications. We used *bwa mem* to map each read to the hg19 reference genome, retaining only uniquely mapped reads. For chimeric reads overlapping *HindIII* cut sites, we only used the 5’ end. We kept reads within 500 bp of the nearest *HindIII* cut site, and removed any intra-chromosomal read pairs within 15 Kb ^71^. Processed data from our eHi-C and external Hi-C were binned into 100 Kb, 40 Kb, and 10 Kb resolution contact matrices for downstream analysis. As shown in ***Figure S1a***, we evaluated Hi-C read summary statistics including: total number of uniquely mapped reads per sample (PCR duplicates removed), total number of intra-chromosomal reads, and total number of informative intra-chromosomal reads which are >15 Kb. For comparison, we include in ***Table S1*** all Hi-C human tissue data available as of mid-2018. Hi-C comparison data included Schmitt et al. ^32^ (14 adult tissues and 7 cell lines) and Won et al. ^17^ (3 paired fetal samples from brain germinal zone and cortical plate). Data quality from this study was comparable to prior studies although our read depth is the highest of any currently available Hi-C dataset from human brain tissue.

We evaluated the reproducibility of the 40 Kb bin resolution eHi-C contact matrices. The biological replicates of each sample showed high reproducibility (Pearson correlation coefficients > 0.96) enabling pooling of all biological replicates for downstream analyses. eHi-C analyses are based on N=3 adult and N=3 fetal cortex samples for chr1-chr22 and N=2 male adult and N=2 male fetal samples for chrX (given that chromatin interactions are distinctive in females).

#### A/B compartments

An output of Hi-C is determination of “A” and “B” compartments corresponding to contiguous regions of active (A) and inactive chromatin (B) which tend to self-associate. A/B compartment analysis was accomplished with an in-house pipeline following initial paper by Lieberman-Aiden et al. ^33^. Hi-C data from our adult cortex, our fetal cortex, fetal germinal zone and cortical plate samples from Won et al. ^17^, and the 21 Hi-C datasets from Schmitt et al. ^32^ were all processed identically. We identified A/B compartments at 100 Kb bin resolution. We applied quantile normalization using the R package “preprocessCore” for batch effect removal across samples. We then applied PCA to the quantile normalized matrix of compartment scores, and graphed samples in PC1 vs. PC2 plots. The PCA analysis shows clear distinctions between human brain and non-brain tissues and cell lines, and developmental stage differences for the brain samples.

#### Topographically associated domains (TADs)

We identified TAD boundary regions using an in-house pipeline to implement the insulation score method, as described in Crane et al. ^72^. Starting from 40 Kb bin resolution raw Hi-C contact matrix we applied HiCNorm ^73^ to obtain the normalized chromatin interaction frequency matrix. Next, each 40 Kb bin, one at a time treated as the anchor bin, obtained its “insulation score” by calculating the sum of normalized interaction frequency between the anchor bin and all bins ±1 Mb of the anchor bin. We further performed quantile normalization on the insulation score across all samples using the R package “preprocessCore”. Finally, we called a 40 Kb bin a TAD boundary region if its insulation score is the minimal in its local neighboring ±1 Mb region.

#### Frequently interacting regions (FIREs)

Following our prior study ^32^, we applied an in-house pipeline to identify FIREs using 40 Kb resolution Hi-C contact matrices for each chromosome. For each bin, we calculated the total number of *cis* (intra-chromosomal 15-200 Kb) interactions. We then applied HiCNormCis ^32^ to remove systematic biases from local genomic features, including effective fragment size, GC content and mappability ^73^. We removed 40 Kb bins in the MHC region. After filtering, 64,222 40 Kb bins remained for analysis. The normalized total number of *cis* intra-chromosomal reads is defined as the FIRE score ^32^. We then performed quantile normalization of FIRE scores across all samples using the R package “preprocessCore” to remove potential batch effects among different samples. We transformed FIRE score using log_2_(FIRE score+1), and transformed FIRE scores to *Z*-scores (mean=0 and standard deviation=1). We designated FIRE bins as 40 Kb regions with a FIRE score one-sided *P*-value < 0.05. FIREs often cluster into contiguous runs of bins termed super FIREs. We used an in-house pipeline to call super FIREs, motivated by super enhancer calling algorithms ^74,75^. The method is also described in our previous study ^32^. For each Hi-C sample, we began with 40 Kb FIRE bins as described above. We then merged consecutive 40 Kb FIRE bins into contiguous FIRE regions, allowing for up to one 40 Kb bin gap. We ranked these contiguous FIRE regions by their cumulative Z-scores, and plotted the ranked FIRE regions as a function of their cumulative Z-score. Finally, we identified the inflection point of such plot, and designated all FIRE regions to the right of the inflection point as super FIREs.

#### Chromatin interactions

We applied a combination of Fit-Hi-C ^76^ with default parameters and our FastHiC ^77,78^ to detect long-range chromatin interactions. Starting from 10 Kb bin resolution raw Hi-C contact matrices, we removed any 10 Kb bin overlapping the ENCODE “blacklist” (uniquely mappable regions that nonetheless yield artificially high signal, ***URLs***) or the MHC region. We then ran the Fit-Hi-C+FastHiC combination caller on all 10 Kb bin pairs ≥20 Kb and ≤2 Mb apart, resulting in a total of 43,222,677 bin pairs. We first applied Fit-Hi-C to generate informative initial values, from which we ran our FastHiC. We applied a stringent Bonferroni correction, and only considered 10 Kb bin pairs with *P* < 0.001/43,222,677 or *P* < 2.31×10^-11^ as statistically significant chromatin interactions. Since we used the six base pair restriction enzyme *HindIII* in eHi-C, it was not possible to directly apply HiCCUPS, another method to detect chromatin interactions (predominantly CTCF-mediated chromatin loops) as it was designed for *in situ* Hi-C data with the four base pair restriction enzyme *MboI*. Our approach is consistent with the long-range chromatin interaction calling algorithm adopted by Won et al. ^17^. Because Fit-Hi-C and FastHiC model the global background, the resulting chromatin interactions identified (i.e., 3D peaks called) are enriched with long-range enhancer-promoter interactions. In contrast, HiCCUPS adopts a local background, and thus the resulting peaks are enriched with CTCF mediated chromatin loops.

### Additional functional genomic data

To aid in the interpretation of our eHi-C data, we generated additional data from human brain samples. ***Table S2*** summarizes the data types and sample developmental stage/brain region. Methods for the data we generated are provided below, and methods for external data can be found in the primary publications.

#### RNA-sequencing

We generated bulk-tissue RNA-sequencing data from nine fetal cortex and nine adult DLPFC (dorsolateral prefrontal cortex) samples. The collection of dorsolateral prefrontal cortex and the psychiatric characterization are detailed in Zhu et al. ^79^. All controls had no neurological disease or severe mental illness. All samples were free from large structural variants (>100 Kb) detectable using Illumina OmniExpress arrays. Genotypic sex matched phenotypic sex for all samples. We extracted total RNA from 25 mg of pulverized tissue per sample using Norgen’s Fatty Tissue RNA Purification Kit (Norgen Biotek, Thorold, ON Canada). Extracted RNA from the cell pellets (each with ~3-6 million cells) using Norgen’s Total RNA extraction kit. RNA concentration was measured using fluorometry (Qubit 2.0 Fluorometer), and RNA quality verified using a microfluidics platform (Bioanalyzer, Agilent Technologies). Barcoded libraries were created (Illumina TruSeq RNA Sample Preparation Kit v4) using 1 μg of total RNA as input. Samples were randomly assigned to bar codes and lanes. Libraries were quantified using fluorometry and equal amounts of all barcoded samples were pooled and sequenced using Illumina HiSeq 2000 (100 bp paired-end). We used the quasi-mapping-based mode of Salmon (version 0.8.2) ^80^ to generate transcript-level counts from RNAseq reads using the GENCODE gene models (v26). We used tximport to generate gene-level counts ^81^ and DESeq2 ^82^ for differential expression analysis. We used the WGCNA R package ^83^ for co-expression network analysis of our RNA-seq data from fetal and adult cortex. We also generated total-stranded RNA-seq from one control DLPFC sample and one fetal cortical sample. The purpose was to enable detection of RNA sequences present in brain but not in GENCODE gene models.

#### Open chromatin

Assay for transposase-accessible chromatin sequencing (ATAC-seq) was used to map chromatin accessibility genome-wide ^84^. This method probes DNA accessibility with hyperactive Tn5 transposase, which inserts sequencing adapters into accessible regions of chromatin. We used tissue from adult DLPFC with no history of psychiatric or neurological disorders (N=137) ^50^. Approximately 20 mg of pulverized brain tissue was used for ATAC-seq. Frozen samples were thawed in 1 ml of nuclear isolation buffer (20 mM Tris-HCL, 50 mM EDTA, 5mM Spermidine, 0.15 mM Spermine, 0.1% mercaptoethanol, 40% glycerol, pH 7.5), inverted for 5 minutes to mix, and samples were filtered through Miracloth to remove larger pieces of tissue. Samples were centrifuged at 1100 x *g* for 10 min at 4°C. The resulting pellet was washed with 50 μl RSB buffer, centrifuged again, and supernatant was removed. The final crude nuclear pellet was re-suspended in transposition reaction mix and libraries prepared for sequencing as described in Buenrostro et al. ^84^. All samples were barcoded, and combined into pools. Each pool contained 8 randomly selected samples (selection balanced by case/control status and sex). Each pool was sequenced on two lanes of an Illumina 2500 or 4000 sequencer (San Diego, CA, USA). Raw fastq files were processed through cutadapt (version 1.2.0) ^85^ to remove adaptors and low-quality reads. cutadapt-filtered reads were aligned to hg19 using Bowtie2 ^70^ using default parameters. In alignment, all reads are treated as single-read sequences, regardless of whether ATAC-seq libraries were sequenced as single-end or paired-end. The aligned bam files were sorted using samtools (version 0.1.18) ^86^, duplicates removed using Picard MarkDuplicates, and then converted to bed format using BedTools (version: v2.17.0) ^87^. ENCODE “blacklist” regions (***URLs***) were removed (i.e., empirically identified regions with artefactual signal in functional genomic experiments). Narrow open chromatin peaks were called from the final bed files using MACS2.

#### ChIP-seq

We generated epigenetic marks using postmortem adult and fetal cortex samples (H3K27ac, H3K4me3, and CTCF). All assays were done using the ChIPmentation protocol ^88^. Brain tissues were crosslinked with 1% formaldehyde at room temperature followed by glycine quenching. To isolate nuclei, tissues were lysed with Lysis buffer I (10 mM HEPES; pH 7.5, 10 mM KCl, 0.1 mM EDTA, 1 mM dithiothreitol, 0.5% Nonidet-40, and protease inhibitor cocktail) for 10 minutes at 4°C. The collected nuclei were then washed with a lysis buffer II (200mM NaCl, 1 mM EDTA pH 8.0, 0.5 mM EGTA pH8.0, 10 mM Tris-Cl pH 8.0 and protease inhibitor cocktail) for 20 minutes at room temperature. The nuclei pellets were resuspended in lysis buffer III (10mM Tris-Cl pH 8.0, 100 mM NaCl, 1 mM EDTA, 0.5 mM EGTA, 0.1% sodium deoxycholate, 0.5% N-lauroylsarcosine, and protease inhibitor cocktail) for sonication. The chromatin was sheared for 10 cycles (15 seconds on and 45 seconds off at constant power 3) on Branson 450 sonifier. For the pulldowns, 20-50 μg of chromatin was used for H3K4me3 (Abcam, ab8580) and H3K27ac (Abcam, ab4729), and 100-150 μg for CTCF (Abcam, ab70303). First, 11 μl of Dynabeads M-280 (Life Technologies, Sheep Anti-Rabbit IgG, Cat# 11204D) was washed three times with 0.5 mg/ml of BSA/PBS on ice and then incubated with each designated antibody for at least 2 hours at 4°C. The bead-antibody complexes were then washed with BSA/PBS. The pulldown was down in binding buffer (1% Trixon-X 100, 0.1% sodium deoxycholate, and protease inhibitor cocktail in 1X TE) by mixing the bead-antibody complexes and chromatin. After pulling down overnight, the bead-antibody-chromatin complexes were washed with RIPA buffer (50mM HEPES pH 8.0, 1% NP-40, 0.7% sodium deoxycholate, 0.5M LiCl, 1mM EDTA, and protease inhibitor cocktail). The bead complexes were then subjected to ChIPmentation by incubating with homemade Tn5 transposase in tagmentation reaction buffer (10mM Tris-Cl pH 8.0 and 5mM MgCl_2_) for 10 minutes at 37°C. To remove free DNA, beads were washed twice with 1x TE on ice. The pulldown DNA was recovered by reversing crosslinks overnight followed by PCRClean DX beads purification. To generate ChIP-seq libraries, PCR was applied to amplify the pulldown DNA with illumina primers. Size selection was then done with PCRClean DX beads to choose fragments ranging from 100-1000 bp. All ChIP-seq libraries were sequenced on Illumina HiSeq2500 platform (50 bp single-end). All ChIPmentation reads were mapped to hg19 of the human genome using Bowtie ^70^. The first 36 bases of each reads were applied for alignment with up to 2 mismatches allowed. To remove duplication, only uniquely mapped reads were kept for further analysis. Peak calling was performed using MACS2 ^89^.

### Comparing Hi-C readouts to human genetic results

We compared the eHi-C readouts to multiple sets of human genetic results selected due to the availability of large GWAS (***Table S2***). We evaluated the connection between Hi-C readouts and these GWAS using partitioned LD score regression ^90,91^. In effect, we estimate the degree to which the SNP-heritability of a trait is enriched in a set of genomic features (e.g., FIREs). Partitioned LD score regression is an extension of LD score regression allowing to estimate whether one or more sets of pre-specified genomic regions are enriched for the SNP-heritability of a trait based on GWAS summary statistics. Briefly, LD score regression ^92^ estimates common-variant SNP heritability by regressing the χ^2^ association statistics from GWAS against LD scores (the sum of r^2^ for each SNP in a reference population). For multigenic traits, SNPs in high LD should on average be more associated with the trait than SNPs in low LD as they are expected to capture more of the genetic effects on the trait. The relationship between χ^2^ statistics and LD scores is directly dependent on the proportion of genetic variance of the trait, which allows estimation of SNP-heritability ^92^. Partitioned LD score regression ^90^ uses the same principle except that SNPs are partitioned into functional categories. If some categories are enriched in causal variants, the relationship between χ^2^ statistics and LD scores should be stronger than for categories with few causal variants. This allows estimation of the degree of enrichment of SNP-heritability in one or more functional categories. LD scores are computed for each category based on the presence of the SNP in the annotation (1 if a SNP is located in the annotation, 0 otherwise). For each annotation of interest (e.g. FIREs), we added SNPs from the “baseline model” located within the genomic coordinates of the regions as an extra annotation (1 if a SNP is in the region, 0 otherwise). For heritability enrichment, we added an extra annotation surrounding the annotation of interest (e.g., FIREs) ±500 bp to prevent upward bias in heritability enrichment estimates (as recommended) ^90^. Significance was assessed using the enrichment P-value, which is not corrected for other genomic annotations. For comparison across tissues, we only added the annotation of interest and used the coefficient Z-score P-value, which is corrected for other genomic annotations (as recommended) ^90^.

### Connecting GWAS results to tissues using bulk-tissue RNA-seq

We obtained summary statistics for multiple human traits that had been the subject of large GWAS (***Table S2***). Our main interests were schizophrenia and intelligence. We obtained bulk tissue RNA-seq gene expression data from GTEx (v7) ^43^. We used partitioned LD score regression ^90^ to test whether the top 10% most specific genes of each tissue ^49^ were enriched in heritability. SNPs located in the top 10% most specific genes (±100 Kb) were added to the baseline model. We then selected the heritability enrichment P-value as a measure of the association with the traits.

### Identifying genes implicated by GWAS

The functional datatypes are listed in ***Table S2***. We focused on adult and fetal cortex. For schizophrenia, for examples, cortex is implicated using orthogonal functional genomic data (bulk tissue mRNA-seq, single-cell RNA-seq, enhancer marks, and open chromatin) ^28,49^-^51^. We selected an inclusive set of protein-coding genes with expression in human brain (GTEx v7, TPM>1 in any brain region or in fetal cortex) ^43,44^.

We selected eQTL SNP-gene pairs from CommonMind frontal cortex, GTEx in any brain region (*q*<0.05), or in fetal cortex ^18,43,44^.

We identified high-confidence chromatin interactions in cortex (*P*<2.31×10^-11^, Bonferroni correction of α=0.001 for 43,222,677 possible 10 Kb interactions that were 20 Kb to 2 Mb apart, excluding any that intersected centromeres, ENCODE blacklist regions, or bins with poor mapping qualities). We identified anchors that overlapped enhancers (E) or promoters (P). E regions were the intersection of: the *HindIII* Hi-C fragment within a 10 Kb anchor, open chromatin in cortex ^50,93,94^, and either H3K27ac peak ^95,96^ or H3K4me3 peak atop a brain-expressed transcript start site. P regions were defined as a brain-expressed transcripts overlapping open chromatin in cortex ^50,93,94^. These were identified separately using functional genomic data from adult and fetal cortex. For adult cortex, there were 370,994 high-confidence chromatin interactions (*P*<2.31×10^-11^) of which 75,531 were regulatory interactions (E-P or P-P). For fetal cortex, there were 969,302 high-confidence chromatin interactions of which 75,246 were regulatory interactions (E-P or P-P). A down-sampling analysis showed that the greater number of high-confidence chromatin interactions we due to greater read depth.

We did not apply statistical prioritization algorithms (e.g., TWAS, co-localization, or credible SNP): these widely used and respected approaches were not aligned with our goal of generating an inclusive set of hypotheses. Some of these methods are restrictive – e.g., only 69 of 18K significant SNPs in CLOZUK were credible SNPs with posterior probabilities >0.9. Moreover, to our knowledge, these methods have not been rigorously benchmarked in high-throughput experimental models, and some methods often make the limiting assumption of a single causal SNP per locus.

We compared our approach to Wang et al. ^19^ who evaluated 142 SCZ GWAS loci from the CLOZUK study ^12^. The original study reported 145 significant loci, and Wang et al. excluded the extended MHC region and two chrX loci. Using their Hi-C and QTL data, they reported connections of SCZ loci to 1,111 genes of which 321 were “high-confidence” (i.e., supported by more than one source of evidence). We used the same set of SCZ results but evaluated protein-coding genes expressed in adult or fetal cortex. The list of 1,111 genes in Wang et al. included 786 protein-coding genes and 325 genes of less certain salience: pseudogenes (106), miscellaneous RNA genes (78), antisense transcripts (71), and lincRNAs (70). Of the 786 protein-coding, autosomal, non-MHC genes from Wang et al., 271 were labelled as high-confidence. We found substantial overlap of our result with Wang et al.: 81.0% (596/736) of the protein-coding genes identified by Wang et al. were also found by us, particularly their “high-confidence” genes (95.9%, 260/271). However, we identified many more genes (1,300 vs 736) mainly due to our far deeper eHi-C data.

### Hi-C features and human evolutionary history

We assessed two main ideas. First, we wished to evaluate whether TADs had any relation to LD blocks. TADs and LD blocks are Kb to Mb sized genomic regions that define key regions of interest in functional genomics and human genetics. We did this by evaluating recombination rates and LD decay centered on TAD boundaries. Second, given the importance of brain FIREs, we assessed whether these regions had evidence of evolutionary selection.

#### TAD boundaries and recombination rates

To examine the relationship between TAD boundaries and recombination rates, we downloaded the HapMap recombination map (***URLs***) ^97^. We divided the genome in 40 Kb bins and excluded regions with <10 SNPs as well as those with poor mappability, GC content, centromeric location, or poor performance in functional genomic assays via the ENCODE blacklist. Each remaining 40kb bin was dichotomized as a TAD boundary or not, based on our TAD calling results. For each bin, we calculated minimal, maximal and median recombination rate. Previous studies have suggested that GC content and SNP density both influence recombination rate ^98,99^. We therefore calculated summary statistics for these two factors and tested their differences in TAD boundaries and non-TAD boundary bins.

To test whether recombination rates differ in TAD boundaries, we took two approaches, one based on multiple regression and that other on resampling to account for GC content and SNP density ^98,99^. The regression approach simultaneously adjusted for GC content and SNP density ^98,99^. The resampling approach also controlled for GC content or SNP density. Specifically, let *n* denote the number of TAD boundary bins. We repeated the following procedure 10,000 times: from the set of TAD non-boundary bins, we created a subset *s*_*i*_, *i = 1,2, … 1,000* of length *n* bins, by sampling bins, with replacement, such that the distribution of GC content (or SNP density) in *s*_*i*_ matched that in the set of TAD non-boundary bins. For each *s*_*i*_, we calculated the median (across the *n* bins) of relevant summary statistics (median, mean and max recombination rates), *t*_*i*_. We thus generated an empirical null distribution based on which empirical p-values were derived for significance testing.

#### TAD boundaries and linkage disequilibrium

To examine the relationship between TAD boundaries and LD, we computed LD r^2^ values for variants in the 1000 Genomes Project ^67^. After removing structural variants ^100^ and SNPs overlapping with small insertions and deletions, we grouped SNPs by whether they reside in a TAD boundary bin, similarly as in the previous recombination rate analysis. We then evaluated LD decay pattern for SNPs in or not in TAD boundary bins.

#### Brain FIREs and ancient positive selective sweep

We first determined whether brain FIREs have evidence of positive selection since human divergence from Neanderthals, as indicated by the top 5% Neanderthal selective sweep scores (NSS, ***URLs***) ^101^. We first compared brain FIRE bins to non-FIRE bins to assess enrichment for NSS scores. Since FIRE and non-FIRE bins differ in a number of other aspects, which may also relate to positive selection since divergence from Neanderthals, we performed Fisher’s exact test as well as performed logistic regression analysis adjusting for GC content.

Since FIREs are dichotomized by thresholding on continuous FIRE scores (***Online Methods***), signals may be diluted by lumping the remaining ~95% all as the “other” non-FIRE category. We therefore also (a) contrasted the extremes (i.e., top versus bottom 5% of FIRE score bins, or FIREs versus DIREs, depleted interacting regions) using logistic regression and (b) regressed on continuous FIRE scores. For covariates in regression models, we assessed GENCODE gene density, CTCF intensity, the histone marks brain H3K27ac and H3K4me3), open chromatin in DLPFC, and enhancer status in hippocampus (***Table S2***). We included only those that were nominally significant at 0.05 level as covariates in testing for FIRE effects.

#### Brain FIREs and extent of population differentiation

Global population differentiation (measured by the fixation index, *F*_*st*_) is informative for soft selective sweeps where adaptive mutations increase in frequency but do not reach fixation ^102,103^. We obtained global *F*_*st*_ scores from the 1000 Genomes Selection Browser (***URLs***) ^104^. For each bin, we calculated summary statistics (including mean and median) for *F*_*st*_ scores for SNPs within the bin.

#### Brain FIREs and signals of recent positive selection

Signatures of classical and recent selective sweeps can be gauged by a number of metrics ^105^-^109^. We focused on the integrated haplotype score (iHS) statistic that accommodates variants that have not reached fixation for signals of recent selection, integrating signals over 1,000 generations ^106,109^. We obtained iHS statistics from the 1000 Genomes Selection Browser (***URLs***) ^104^. For each bin, we calculated summary statistics (including mean and median) for iHS scores for SNPs within the bin.

#### Brain FIREs and extremely rare variant frequency

We assessed mutation tolerance for extremely rare variant frequency by comparing singleton and doubleton density based on TOPMed freeze 5b (***URLs***, N=62,784 whole genome sequences).

## URLs

1000 Genomes Selection Browser, http://hsb.upf.edu

ENCODE “blacklist”, http://mitra.stanford.edu/kundaje/akundaje/release/blacklists/hg19-human

FUMA, http://fuma.ctglab.nl/gene2func

gnomAD, http://gnomad.broadinstitute.org

GREAT, http://bejerano.stanford.edu/great

HapMap recombination map, ftp://ftp.ncbi.nlm.nih.gov/hapmap/recombination/2011-01_phaseII_B37

HUGIn, http://yunliweb.its.unc.edu/hugin

Neanderthal selection, ftp://hgdownload.soe.ucsc.edu/goldenPath/hg19/database/ntSssTop5p.txt.gz

NHGRI/EBI GWAS catalog, https://www.ebi.ac.uk/gwas

PsychENCODE portal, https://www.synapse.org//#!Synapse:syn4921369/wiki/235539

Psychiatric Genomics Consortium, http://www.med.unc.edu/pgc/results-and-downloads

SALMON, https://combine-lab.github.io/salmon

TOPMed, https://www.nhlbiwgs.org

## Acknowledgements & Funding

PFS gratefully acknowledges support from the Swedish Research Council (Vetenskapsrådet, award D0886501) and the UNC Departments of Genetics and Psychiatry. MH and IJ were supported by NIH grant U54 DK107977. CC, YY, JSM, and YL are supported by NIH grant R01 HL129132 (awarded to YL). PGR was supported by NIMH grant K01 MH109772. LL, XL, YanL, and FJ are funded by R01 HG009658 (awarded to FJ). P.R.J. was funded by the Sophia Foundation for scientific research (grant nr: S14-27). The primary schizophrenia GWAS data were generated with support from Medical Research Council (MRC) Centre (MR/L010305/1), Program Grant (G0800509) and Project Grant (MR/L011794/1), and funding from the European Union’s Seventh Framework Programme for research, technological development and demonstration under grant agreement n° 279227 (CRESTAR Consortium). The authors acknowledge contributions by Drs. Herbert Meltzer and Bryan Roth, the assistance of Lesa Dieter, and the support of the families of the deceased and the staff of the Cuyahoga County Medical Examiner’s Office, Cleveland, Ohio. The tissue collection and psychiatric assessment was supported by NIGMS COBRE Center for Psychiatric Neuroscience (P30 GM103328, awarded to CAS). We thank the NIH NeuroBioBank for providing the fetal brain samples used in this study. The Genotype-Tissue Expression (GTEx) Project was supported by the Common Fund of the Office of the Director of the National Institutes of Health, and by NCI, NHGRI, NHLBI, NIDA, NIMH, and NINDS. The PsychENCODE Consortium was supported by: U01MH103392, U01MH103365, U01MH103346, U01MH103340, U01MH103339, R21MH109956, R21MH105881, R21MH105853, R21MH103877, R21MH102791, R01MH111721, R01MH110928, R01MH110927, R01MH110926, R01MH110921, R01MH110920, R01MH110905, R01MH109715, R01MH109677, R01MH105898, R01MH105898, R01MH094714, and P50MH106934 awarded to: Schahram Akbarian (Icahn School of Medicine at Mount Sinai), Gregory Crawford (Duke University), Stella Dracheva (Icahn School of Medicine at Mount Sinai), Peggy Farnham (University of Southern California), Mark Gerstein (Yale University), Daniel Geschwind (University of California, Los Angeles), Fernando Goes (Johns Hopkins University), Thomas M. Hyde (Lieber Institute for Brain Development), Andrew Jaffe (Lieber Institute for Brain Development), James A. Knowles (University of Southern California), Chunyu Liu (SUNY Upstate Medical University), Dalila Pinto (Icahn School of Medicine at Mount Sinai), Panos Roussos (Icahn School of Medicine at Mount Sinai), Stephan Sanders (University of California, San Francisco), Nenad Sestan (Yale University), Pamela Sklar (Icahn School of Medicine at Mount Sinai), Matthew State (University of California, San Francisco), Patrick Sullivan (University of North Carolina), Flora Vaccarino (Yale University), Daniel Weinberger (Lieber Institute for Brain Development), Sherman Weissman (Yale University), Kevin White (University of Chicago), Jeremy Willsey (University of California, San Francisco), and Peter Zandi (Johns Hopkins University). CMB acknowledges funding from Swedish Research Council (Vetenskapsrådet Dnr: 538-2013-8864) and the Klarman Family Foundation.

## Supplemental figure legends

**Figure S1.**
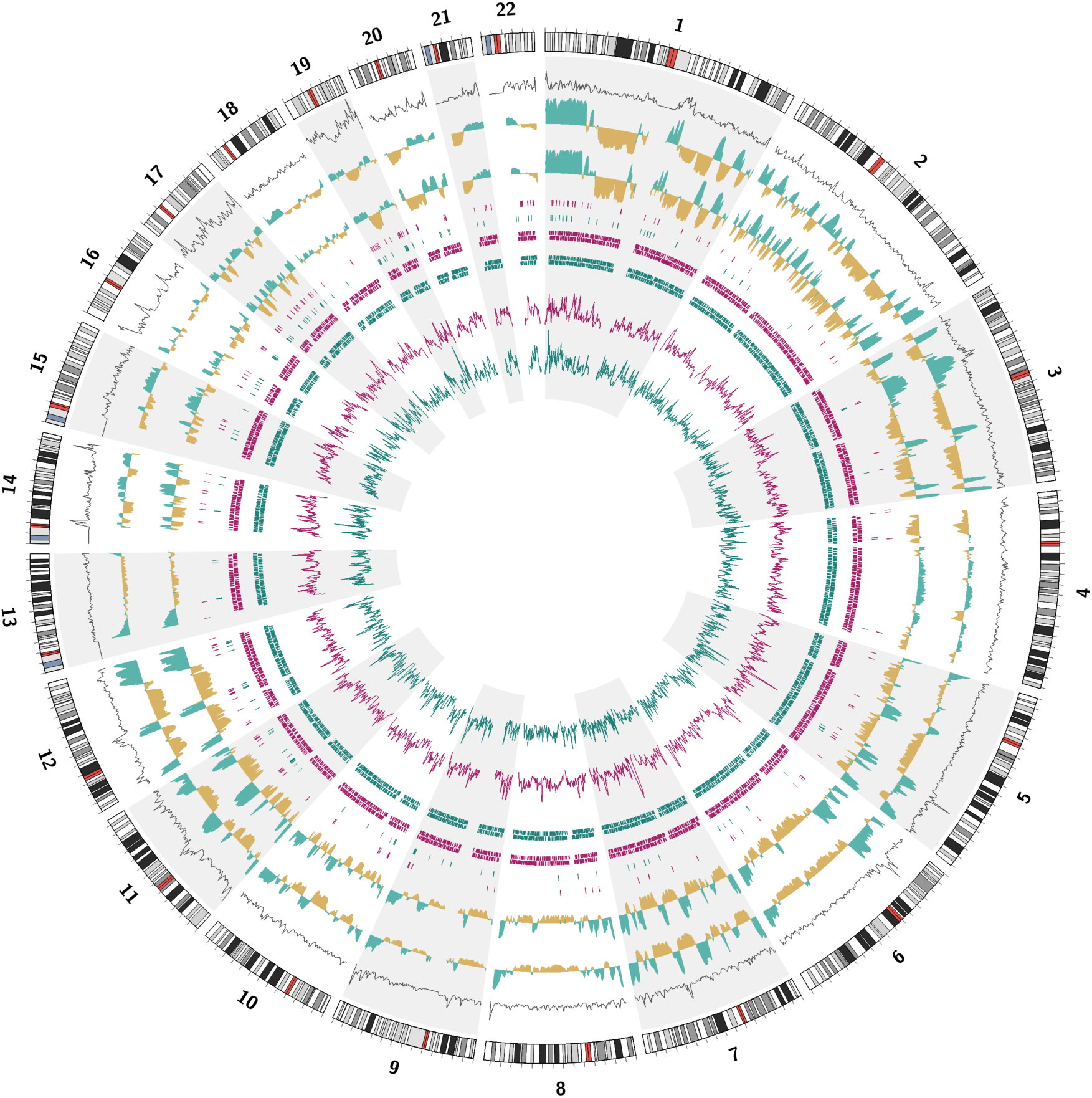
Circos plot summarizing the adult and fetal brain Hi-C readouts in the autosomal genome. From the outside inward, the tracks are: (a) ideogram (chr1ptel to 22qtel in clockwise direction); (b) gene density per Mb; 1 Mb A (green) and B (yellow) compartment in (c) fetal brain and (d) adult brain; locations of super FIREs in (e) fetal brain (red) and (f) adult brain (green); TAD boundary locations in (g) fetal brain and (h) adult brain; and chromatin interaction density per Mb in (i) fetal brain and (j) adult brain.

**Figure S2.**
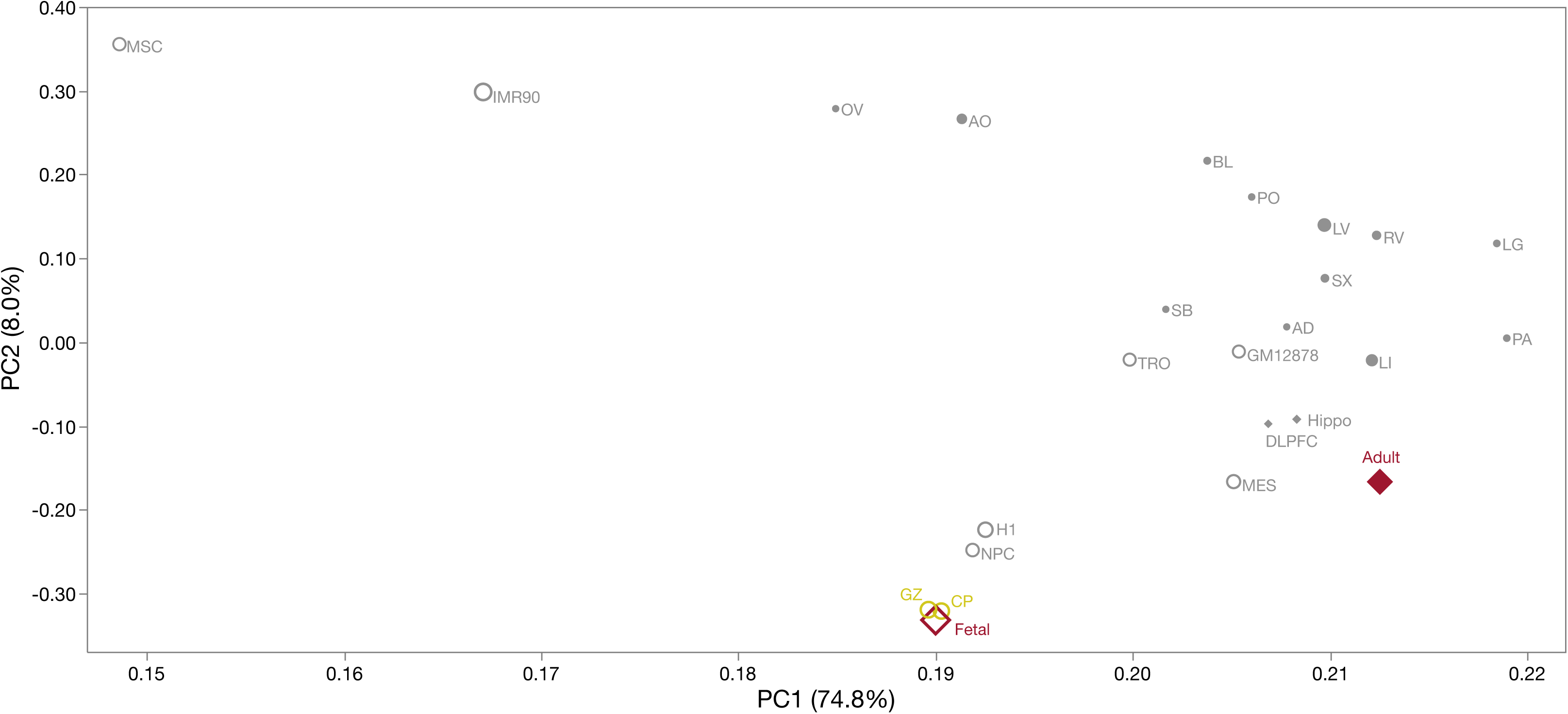
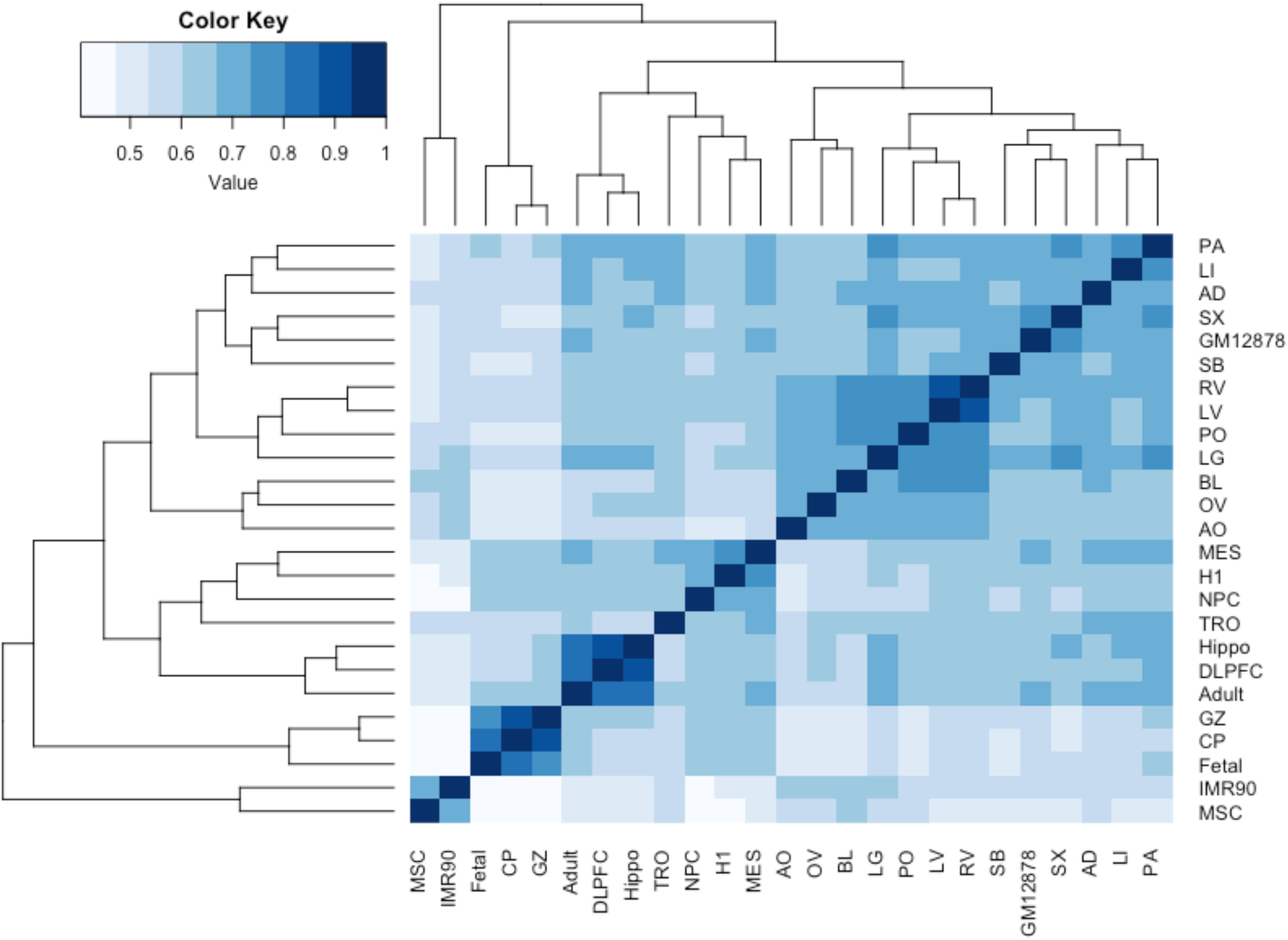
Figure S2a: Results of PCA on Hi-C compartment scores (100 Kb bins) from 25 Hi-C datasets. X-axis is PC1 (75% of variance) and Y-axis is PC2 (8.0% of variance). The fetal samples clustered together as did the adult brain samples. Point sizes are proportional to the numbers of informative *cis*-reads. Filled red diamond is our adult anterior temporal cortex dataset (Adult) and the open red diamond is our fetal cortex dataset (Fetal). Yellow open circles are fetal brain germinal zone (GZ) and cortical plate datasets (CP). Filled grey circles show 14 human tissues and 7 cell lines (see Figure 1a legend for abbreviations). Figure S2b: Clustered heat map of Jaccard index values describing the degree of overlap of compartments across Hi-C datasets.

**Figure S3.**
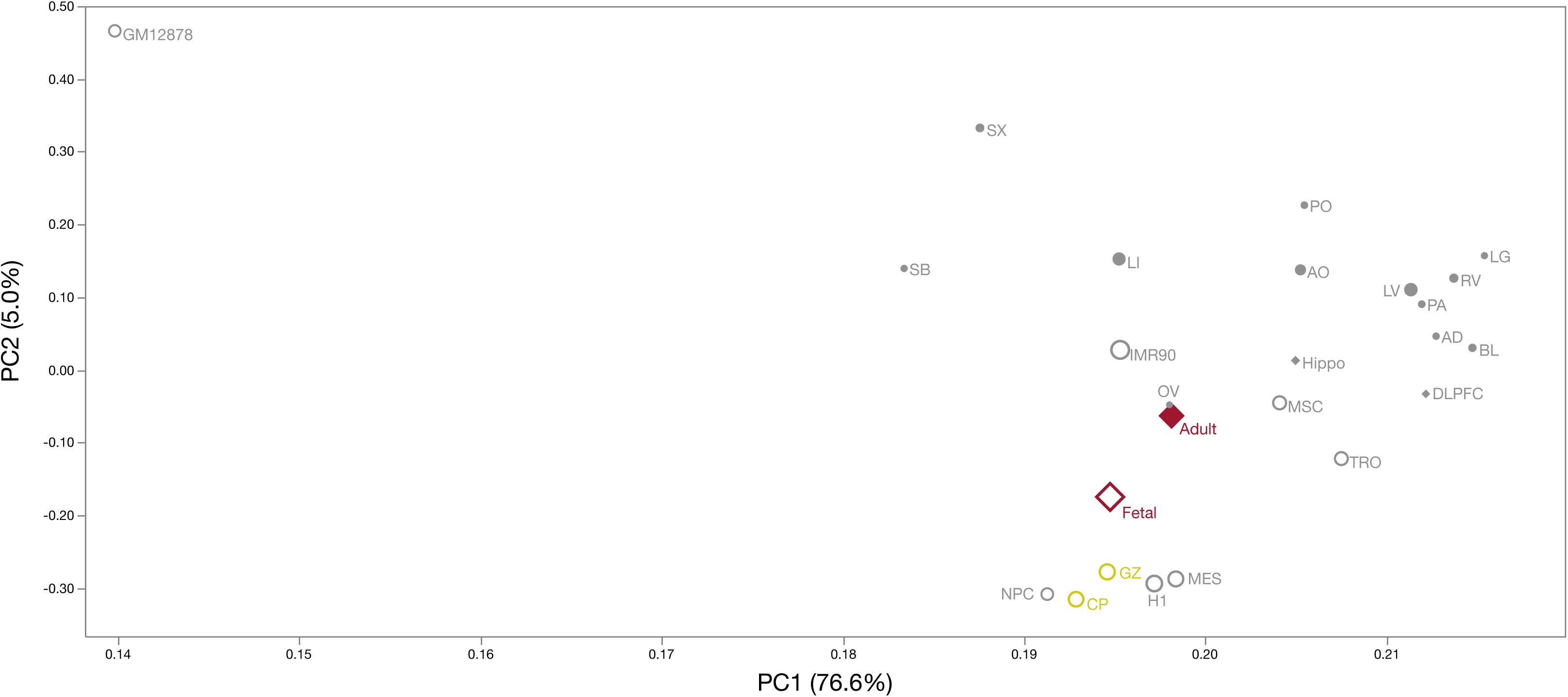
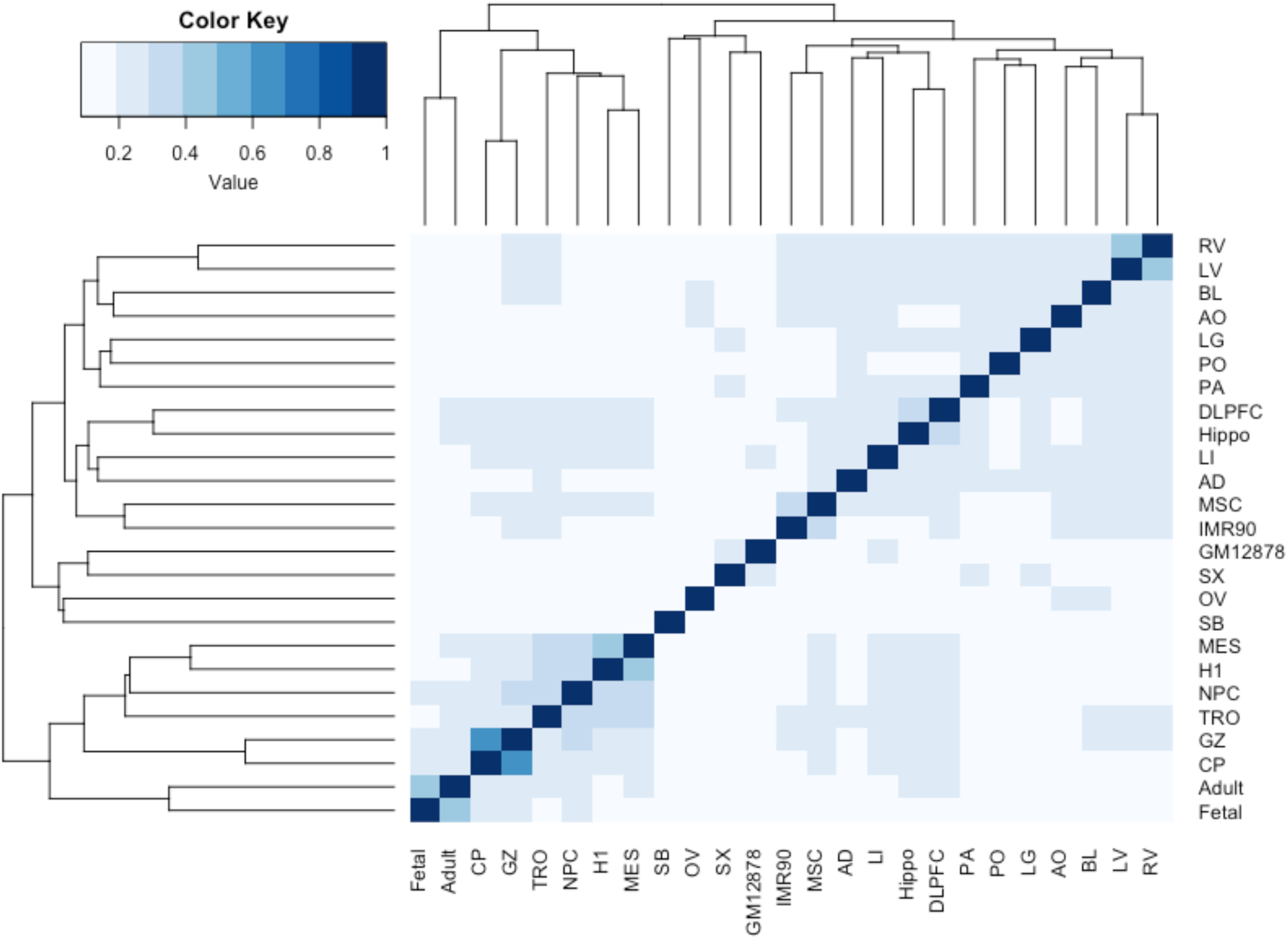
Figure S3a: Results of PCA on Hi-C TAD boundary scores (40 Kb bins) from 25 Hi-C datasets. X-axis is PC1 (77% of variance) and Y-axis is PC2 (5.0% of variance). All brain samples clustered on the dominant PC1. Point sizes are proportional to the numbers of informative *cis*-reads. Filled red diamond is our adult anterior temporal cortex dataset (Adult) and the open red diamond is our fetal cortex dataset (Fetal). Yellow open circles are fetal brain germinal zone (GZ) and cortical plate datasets (CP). Filled grey circles show 14 human tissues and 7 cell lines (see Figure 1a legend for abbreviations). Figure S3b: Clustered heat map of Jaccard index values describing the degree of overlap of TAD boundaries across Hi-C datasets.

**Figure S4.**
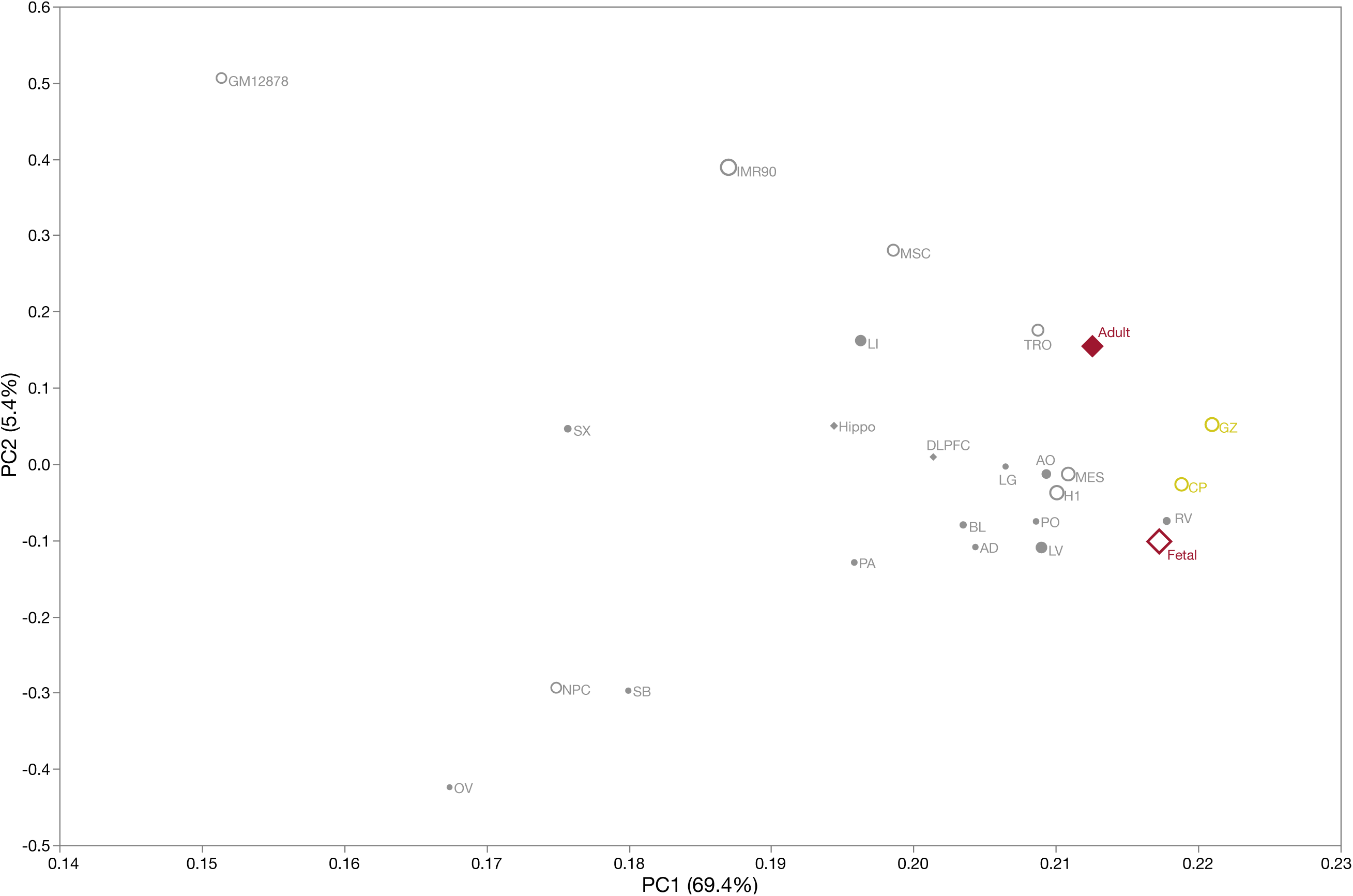
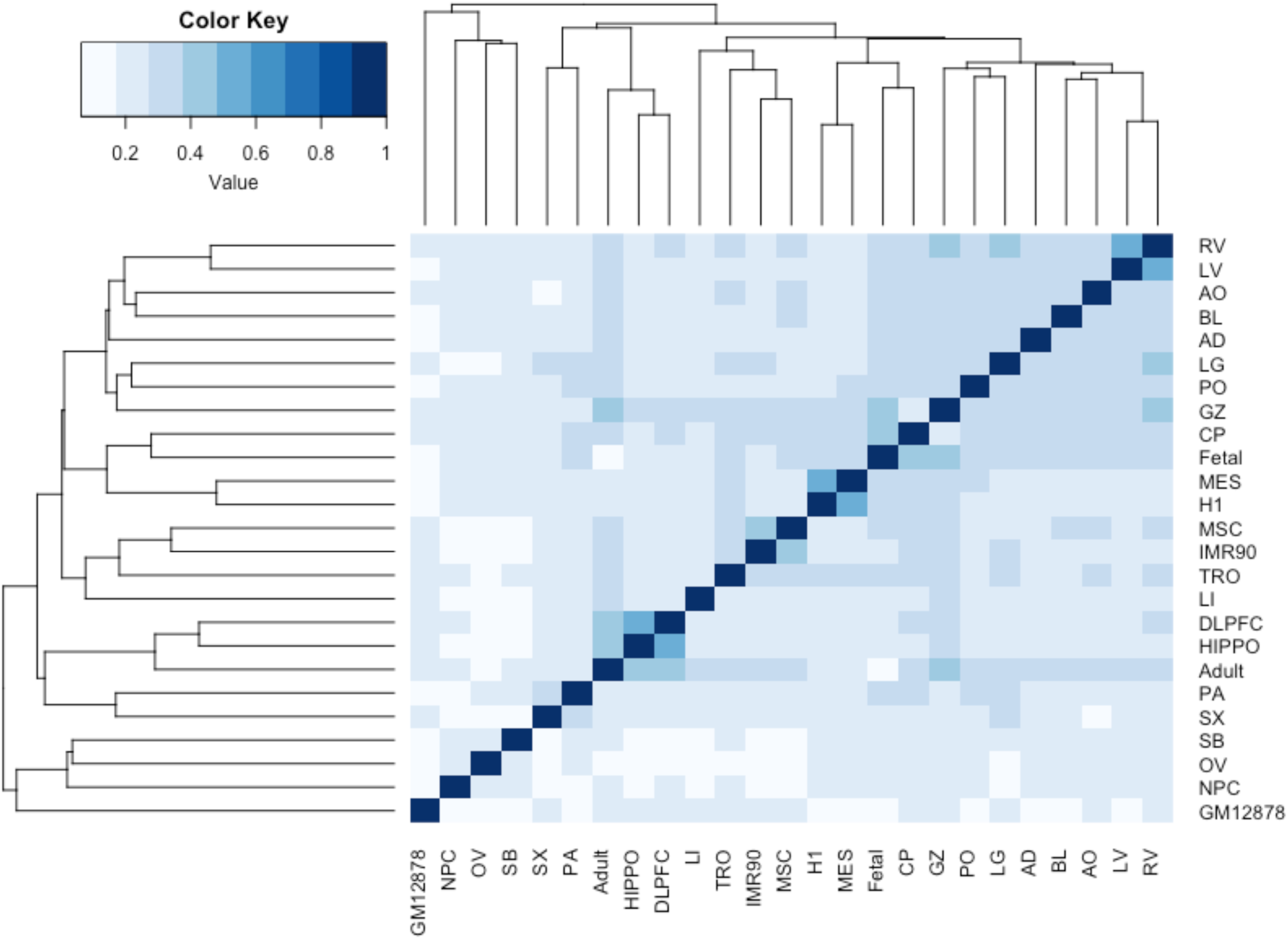
Figure S4a: Results of principal components analysis (PCA) on Hi-C FIRE scores (40 Kb bins) from 25 Hi-C datasets. X-axis is PC1 (69% of variance) and Y-axis PC2 which captured far less variance (5.4%). All brain samples had high scores on PC1. Point sizes are proportional to the numbers of informative *cis*-reads. Filled red diamond is our adult anterior temporal cortex dataset (Adult) and the open red diamond is our fetal cortex dataset (Fetal). Yellow open circles are fetal brain germinal zone (GZ) and cortical plate datasets (CP). Filled grey circles show 14 human tissues and 7 cell lines (see Figure 1a legend for abbreviations). Figure S4b: Clustered heat map of Jaccard index values describing the degree of overlap of FIREs across Hi-C datasets.

**Figure S5.**
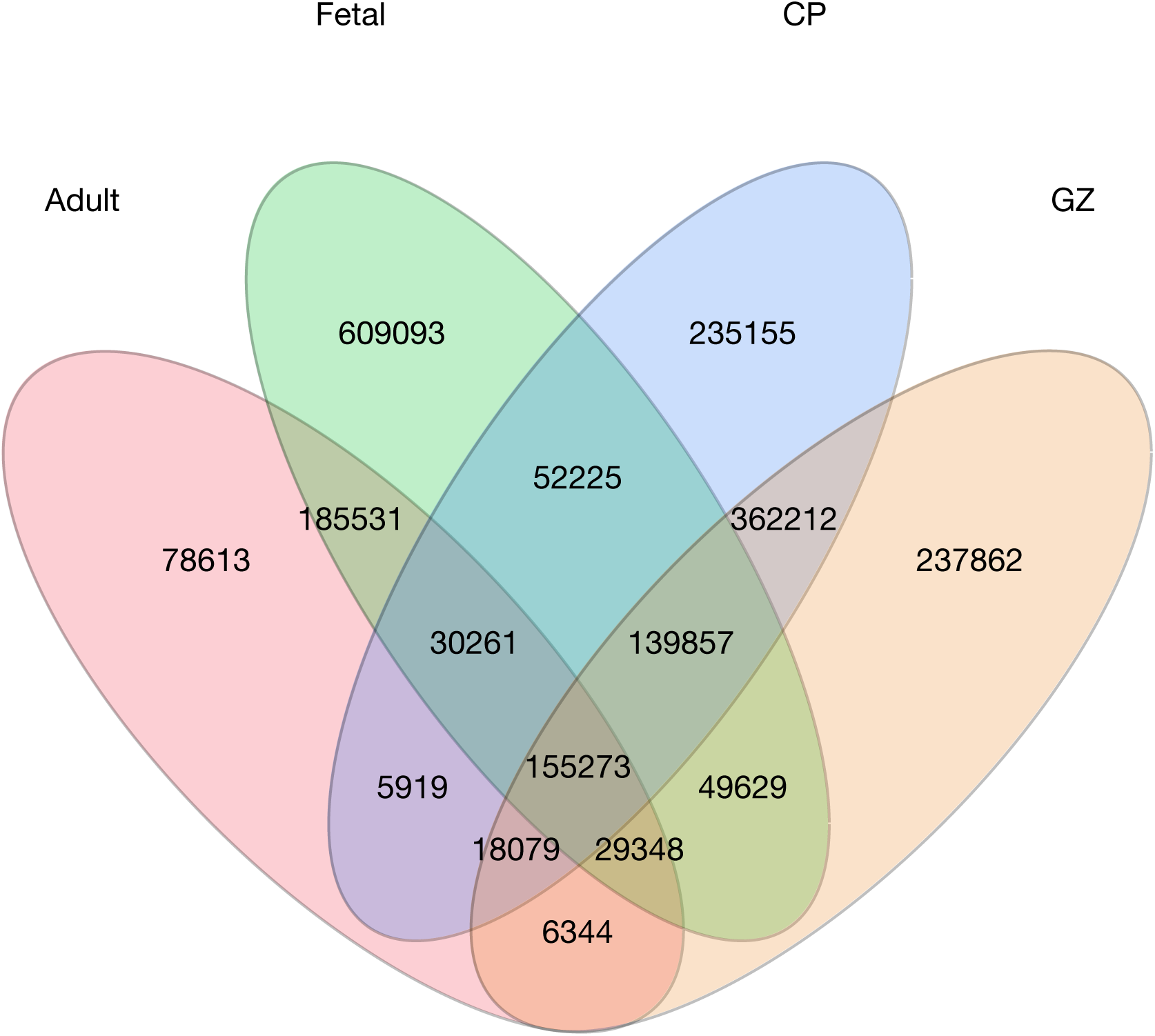
Venn diagram of chromatin interactions in our Adult and Fetal eHi-C datasets and fetal cortical plate (CP) and germinal zone (GZ). Chromatin interactions are between 10 Kb bins that were ≥20 Kb apart and ≤2 Mb apart (i.e., *in cis*). We evaluated these four brain Hi-C datasets because of relatively high read depths (***Figure 1a***). At nominal significance (Bonferroni adjusted *P*<0.05), we identified 2,195,401 chromatin interactions in any of these four brain Hi-C datasets. Fetal cortex had greatest number of chromatin interactions (1.25 million), somewhat more than CP and GZ (0.998 million each), and more than adult cortex (0.509 million).

**Figure S6.**
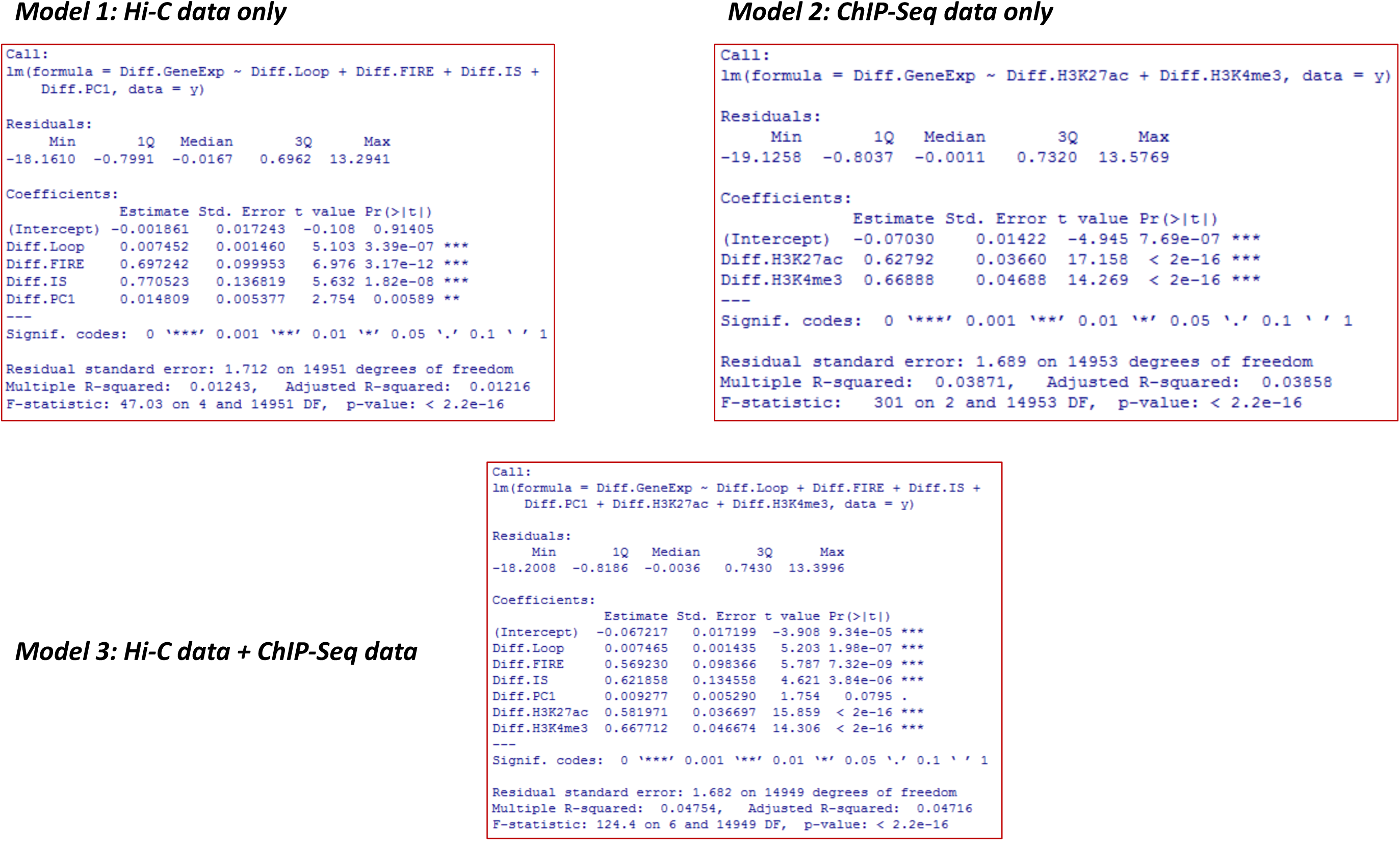
Multivariable models used to evaluate the importance of chromatin interactome on gene expression in fetal and adult brain. A/B compartments, FIREs, and chromatin interactions were significant and orthogonal predictors of gene expression (Model 3, *R*^*^2^*^ 0.0475).

**Figure S7.**
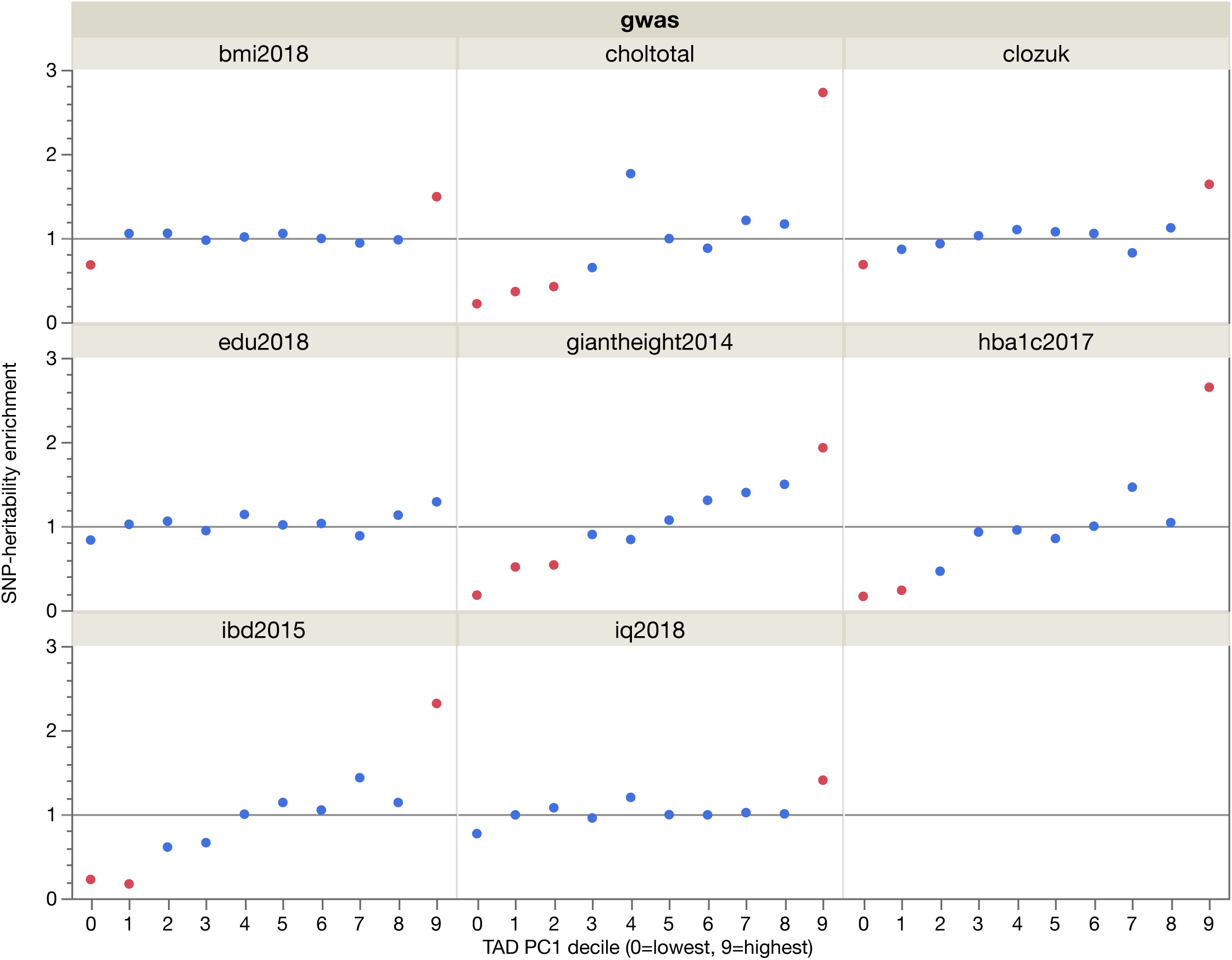
Relationship between TAD PC1 deciles and genome-wide significant findings for complex human disorders and traits. The highest PC1 decile was significant enriched for SNP-heritability while the lowest decile was significantly depleted for SNP-heritability for diverse human traits and disorders. bmi2018 = Body mass index GWAS (Yengo 2018); choltotal = Total cholesterol GWAS (Willer 2013); clozuk = Schizophrenia GWAS (Pardiñas 2018); edu2018 = Educational attainment GWAS (Lee 2018); giantheight2014 = height (GIANT 2014); hba1c2017 = Hemoglobin A1c GWAS (Wheeler 2017); ibd2015 = inflammatory bowel disese (2015); and iq2018 = Intelligence GWAS (Savage 2018).

**Figure S8.**
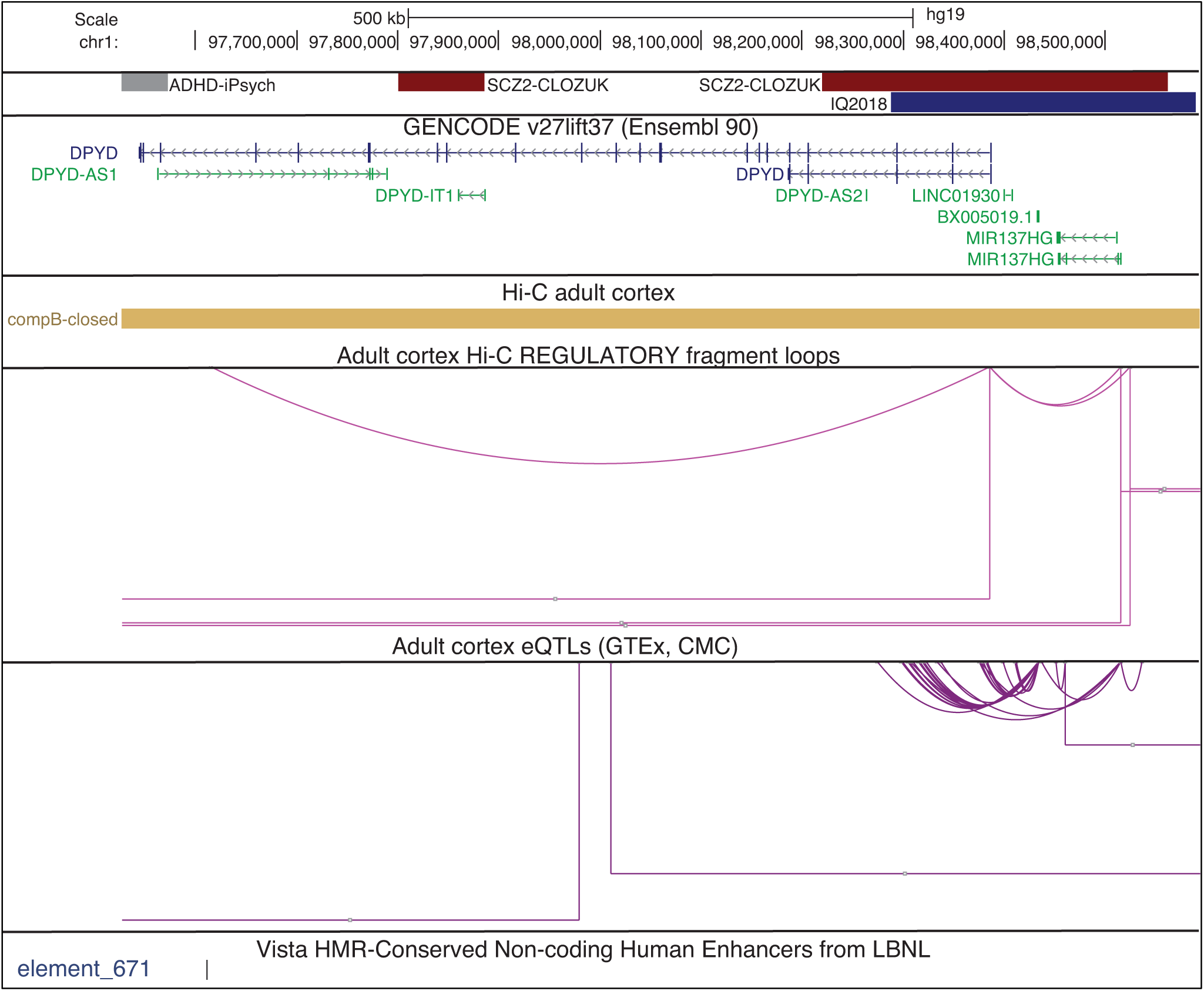
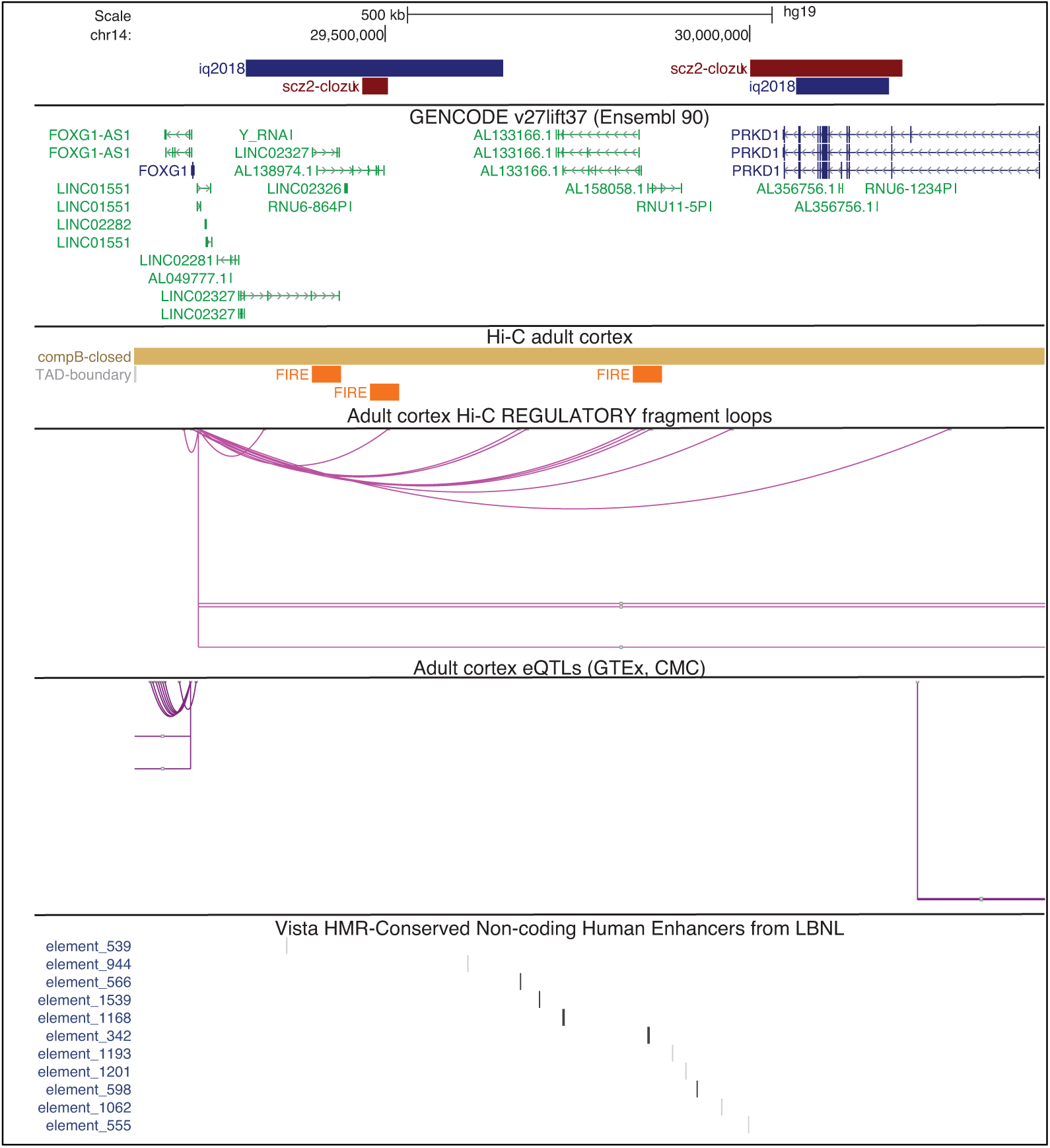
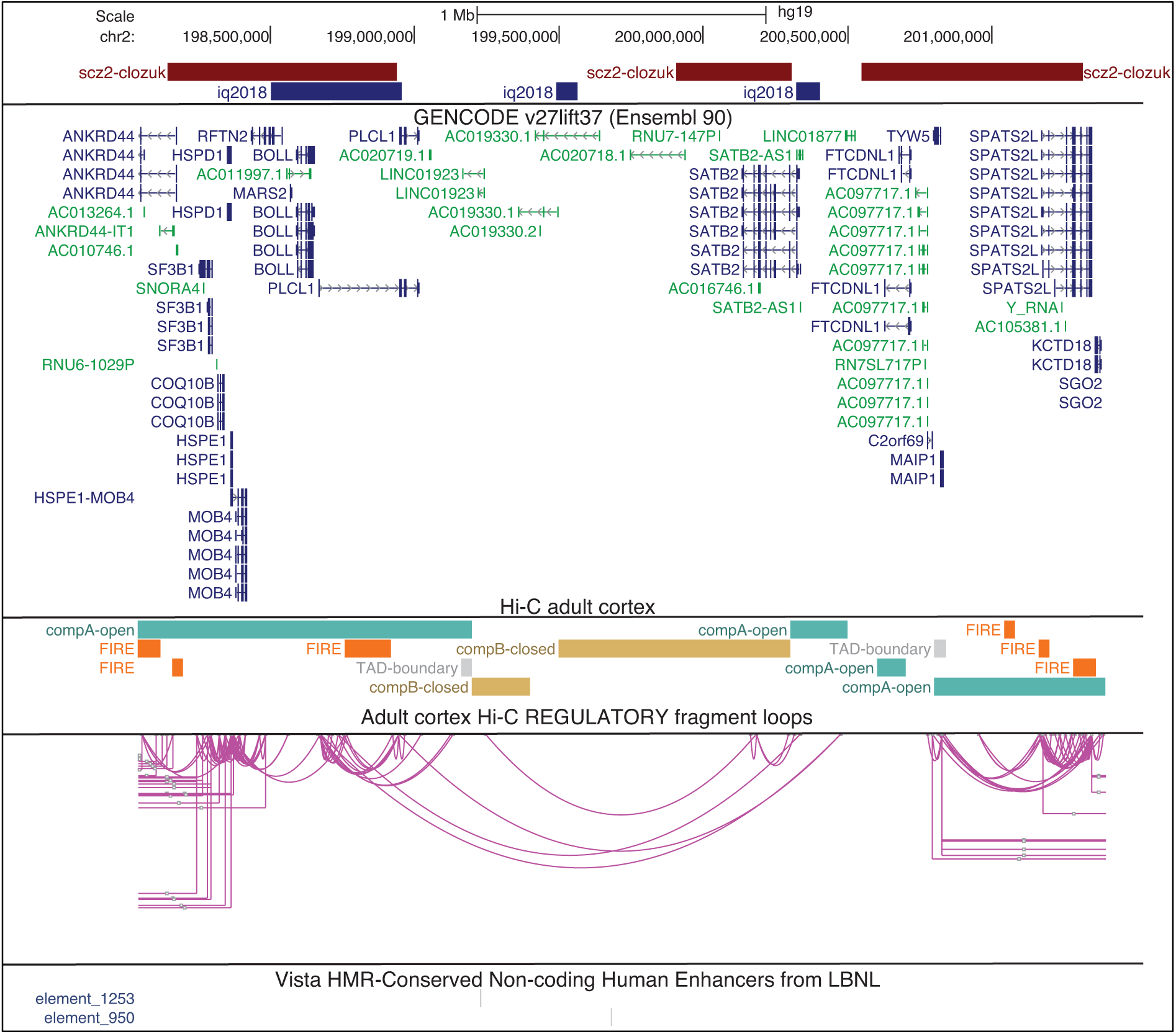
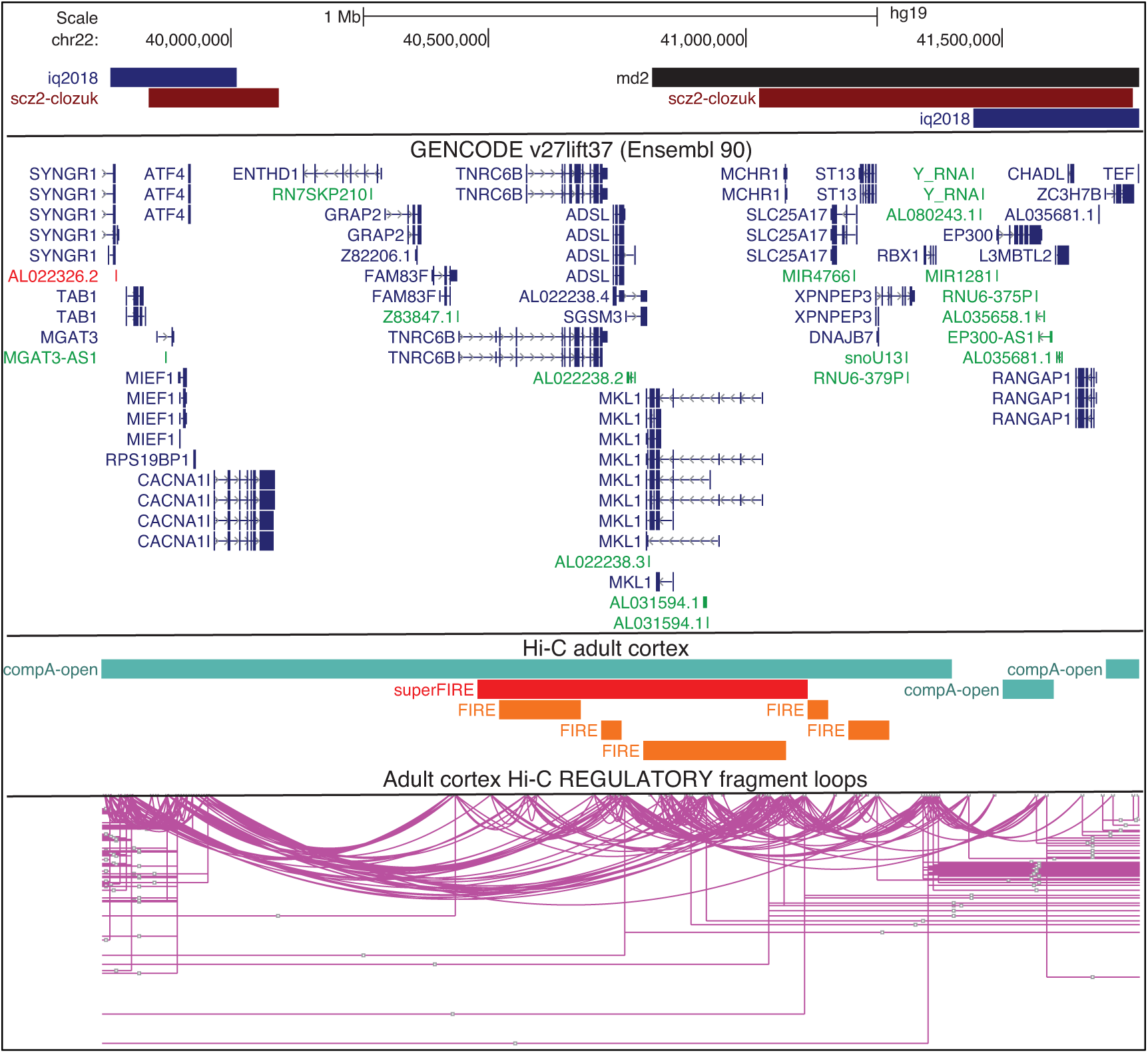
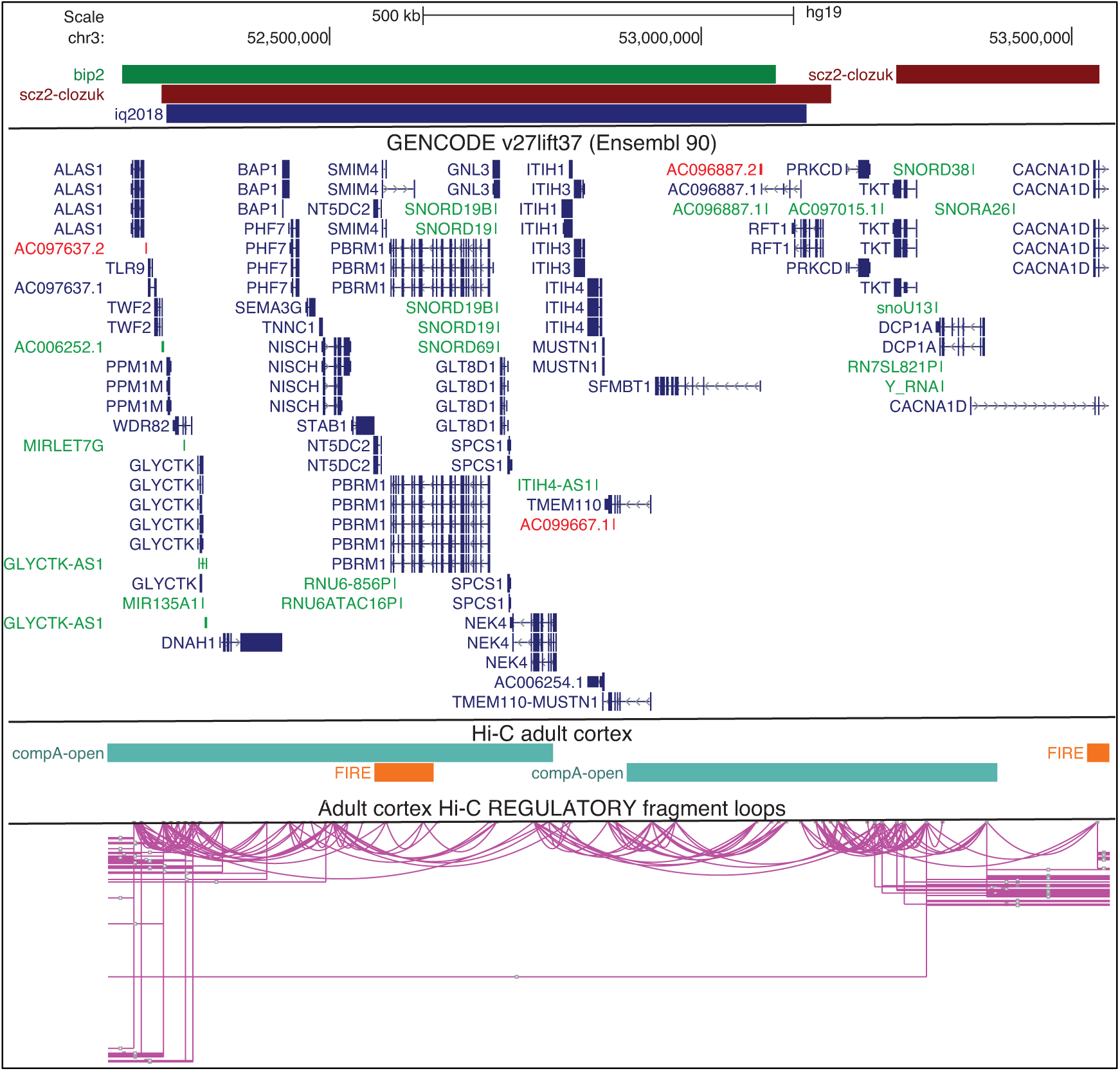
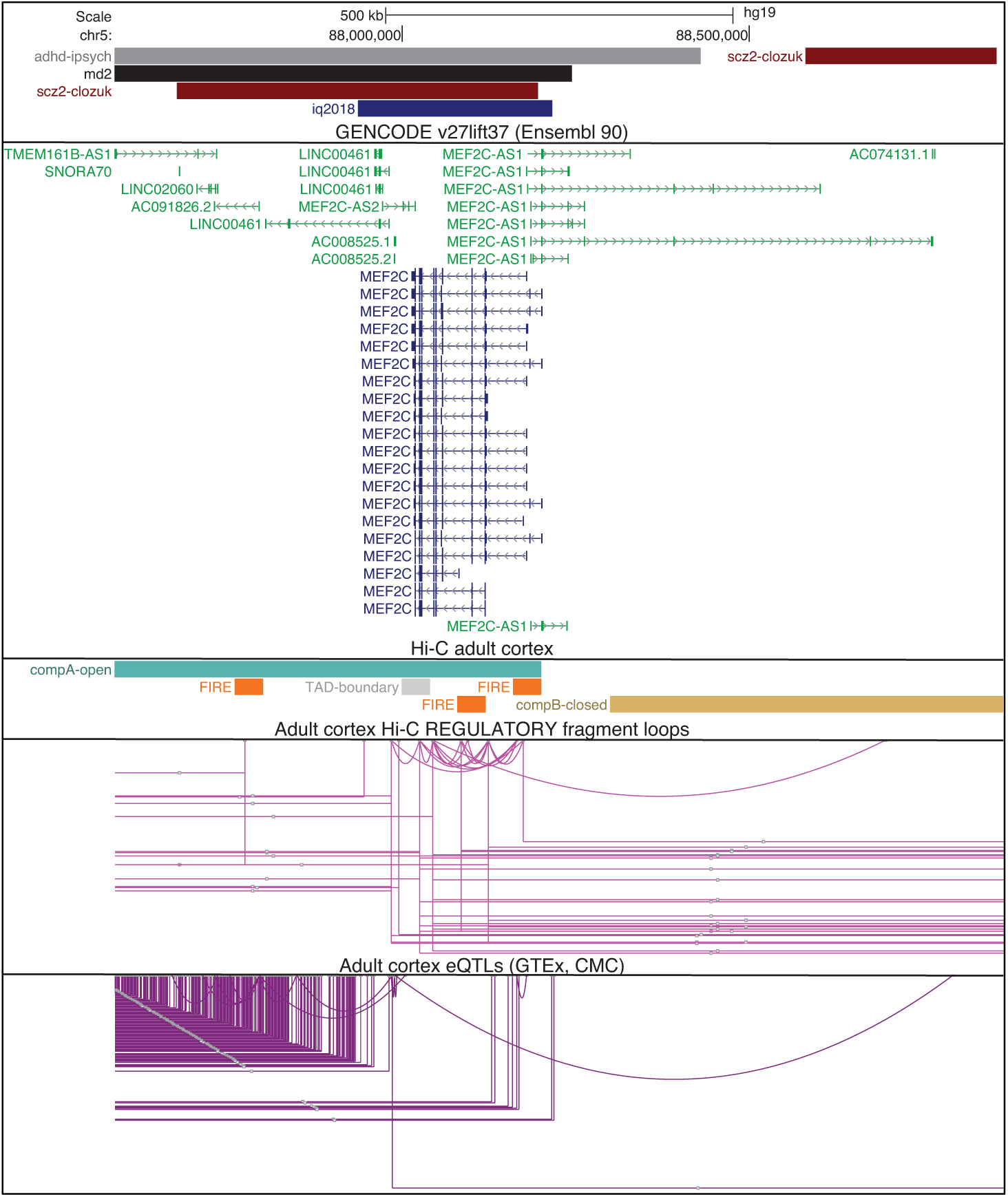
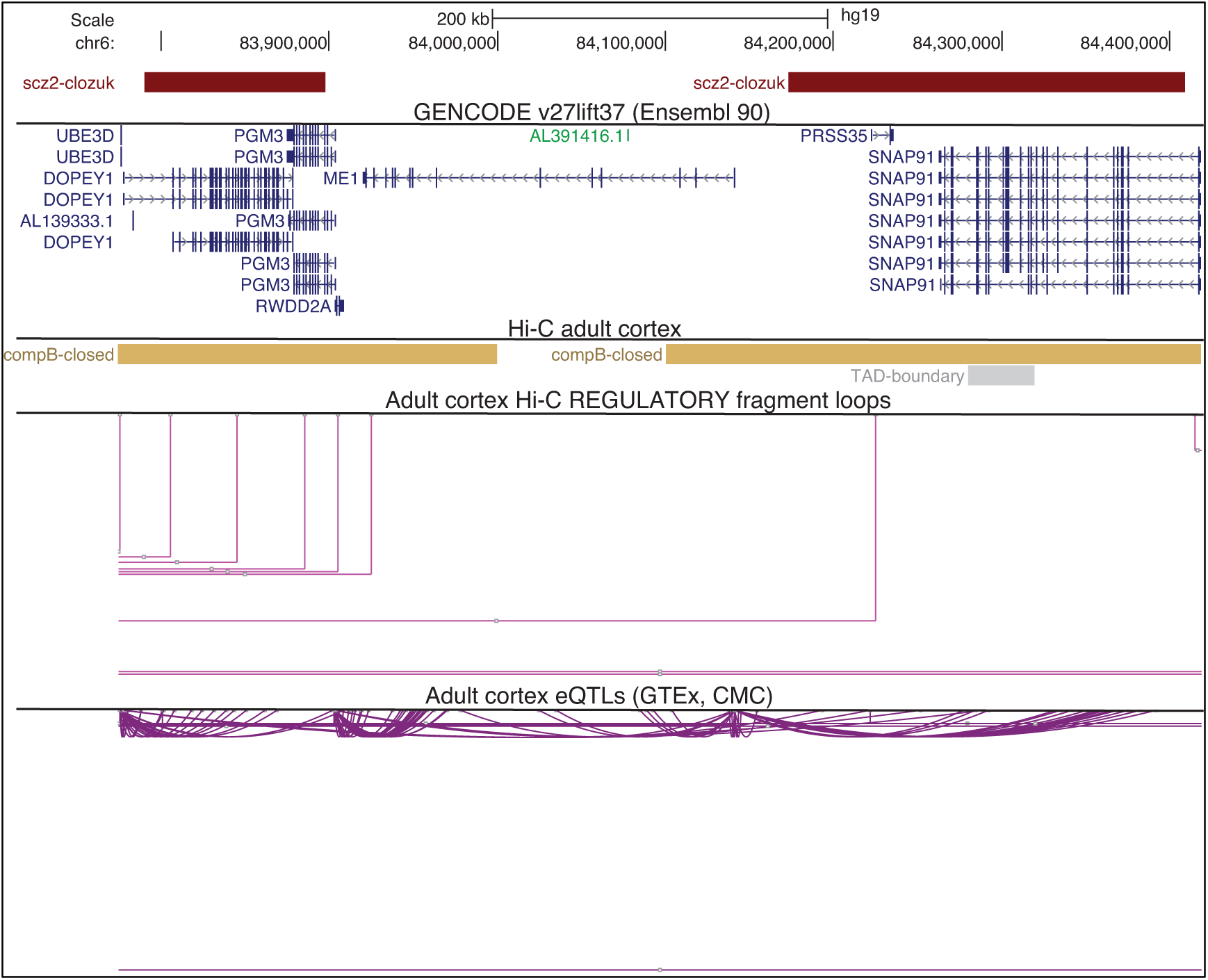
Representative loci with evidence of high-confidence regulatory chromatin interactions for schizophrenia GWAS. UCSC browser tracks (top to bottom): GWAS clumps; GENCODE gene model; adult cortex regulatory chromatin interactions (HCRCI); adult cortex eQTLs; VISTA non-coding human enhancers from LBNL (if present). Figure S8a: *DPYD* (chr1:98,220,320-98,562,260) is a representative example of a single-gene locus. Figure S8b: example of HCRCI bridging two independent loci (chr14:30,000,405-30,208,630 and chr14:29,466,667-29,506,667). Figure S8c: *SATB2* is in chr2:199,908,378-200,305,460 that connects via multiple HCRCI to another locus chr2:198,146,381-198,940,251. Figure S8d: representative example of complex, multigenic loci with dense patterns of HCRCI (chr22:41,027,819-41,753,603 and chr22:39,840,130-40,091,818). Figure S8e: example of a SCZ locus (chr3:52,273,421-53,175,017) that overlaps with loci for bipolar disorder and intelligence. Figure S8f: *MEF2C* (chr5:87,676,693-88,195,380). Figure S8g: schizophrenia loci on chr6:84,173,028-84,409,255 and chr6:83,789,798-83,897,565.

**Figure S9.**
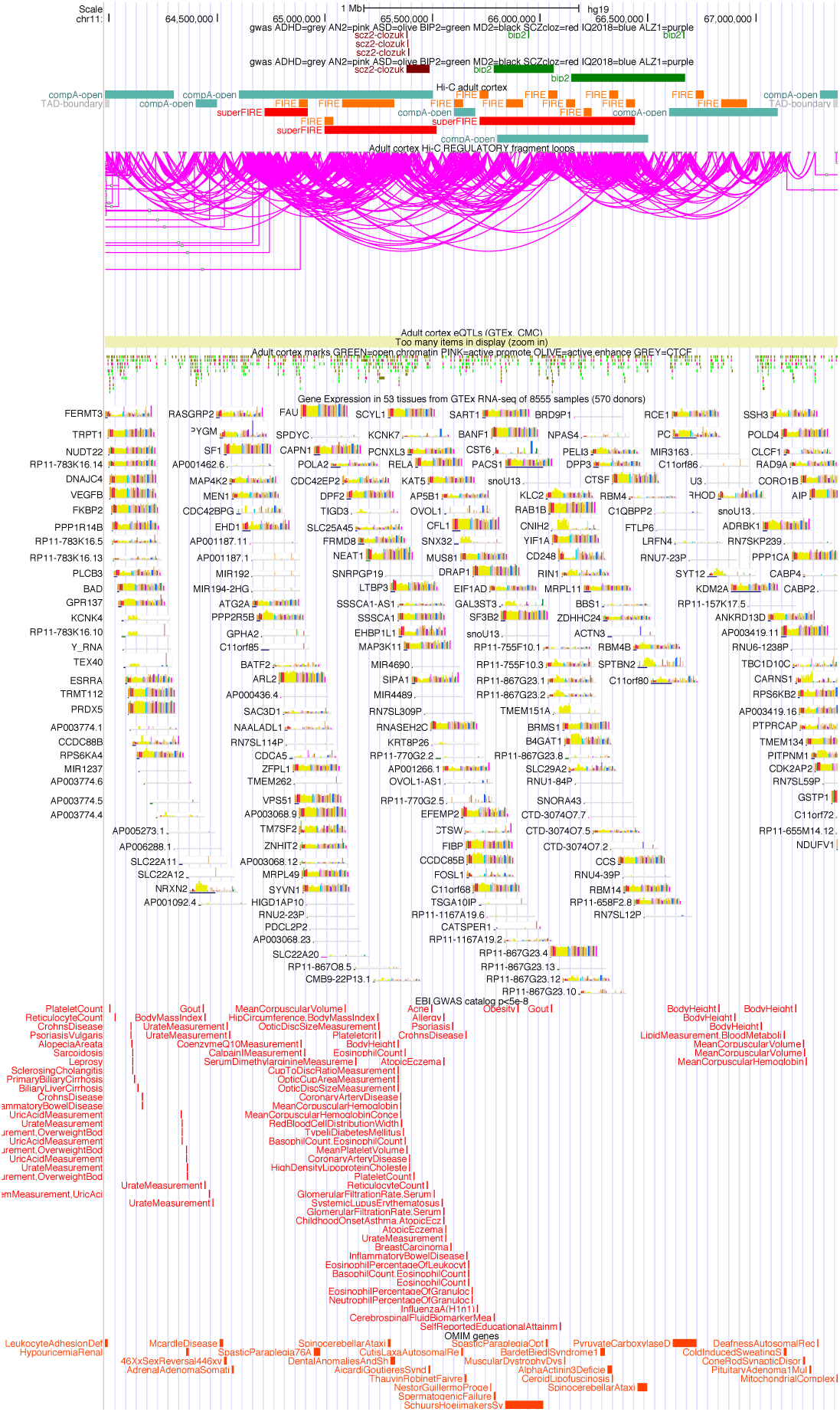
The TAD with the highest value for PC1 (chr11:63980001-67380000). From the top, the tracks show: significant SNP associations, GWAS loci, adult cortex Hi-C readouts, adult cortex HCRCI, adult cortex eQTLs (too many to display), gene expression from GTEx (yellow is for brain tissue), significant SNPs from the GWAS catalog, and OMIM entries. This TAD is dense with genes expressed at high levels across the body and has many disease connections.

